# Scaling genome annotation across the eukaryotic tree of life with OrionGeno

**DOI:** 10.64898/2026.04.26.720859

**Authors:** Lin Liu, Xudong Cai, Shengfu Wang, Yuan Deng, Yiwen Wu, Youliang Pan, Jieyu Wang, Chao Zhang, Haopeng Xia, Nongzhang Tan, Kui Su, Yang Liu, Xuping Zhou, Longqi Liu, Tong Wei, Yong Zhang, Qiye Li, Yuxiang Li, Peng Yin, Xun Xu

## Abstract

The rapid expansion of eukaryotic genome sequencing has created an urgent demand for accurate and scalable genome annotation. Existing *ab initio* methods often struggle to reconstruct complex gene architectures and generalize across distant lineages, limiting their use for large-scale annotation. Here we present OrionGeno, a phylogeny-aware deep learning model for end-to-end eukaryotic genome annotation. OrionGeno integrates phylogenetic context, long-range sequence modeling and joint prediction of gene structures and repetitive elements to annotate exons, introns, untranslated regions and repeats directly from genomic sequences. Applied to chromosome-level eukaryotic genomes from NCBI that lack annotations, OrionGeno generates annotations for more than 5,300 genomes, substantially expanding public annotation resources. Across diverse eukaryotic lineages, OrionGeno outperforms state-of-the-art methods at the exon, gene, protein-sequence, and protein-structural levels. It also identifies candidate protein-coding loci absent from reference protein-coding annotations in well-curated genomes. Together with a web platform and integrated annotation database, OrionGeno provides a scalable and accessible framework for translating genome assemblies into functional biological resources and supporting large-scale biodiversity initiatives such as the Earth BioGenome Project.

## Introduction

The pace of eukaryotic genome sequencing is accelerating rapidly. The Earth BioGenome Project (EBP) is expected to release over 3000 new genomes per month, paving the way for sequencing ∼1.5 million named species of Eukaryota^1,2^. However, accurate gene annotation remains a major bottleneck, with ∼50% of reference genomes in NCBI still lacking annotation. Classic *ab initio* methods, such as Augustus^3,4^, rely heavily on hidden Markov models—an inflexible and “shallow” learning framework that struggles to capture complex exon-intron architectures in DNA sequences. While evidence-driven annotation pipelines^5–7^ can, in principle, identify genes, their accuracy is significantly constrained for species with limited gene expression data or high-quality homologous evidence from closely related organisms. These pipelines are also computationally cumbersome and difficult to deploy at large scale. Consequently, there is a pressing need for a scalable, accurate, and user-friendly tool for genome annotation. Recent deep learning-based models have advanced *ab initio* gene prediction^8–13^, but limitations remain. Fully supervised learning approaches, including Tiberius^8^, Helixer^11^, and ANNEVO^13^, rely on training multiple lightweight models for specific clades, which can improve performance within represented lineages but limits their generalization to novel or under-studied genomes. By contrast, large genomic language model-inspired methods like SegmentNT^12^ use conservation-aware representations pre-trained on genomes from thousands of species, yet they still lag behind specialized tools in gene annotation accuracy^14^. Although effective in certain contexts, these approaches remain limited in their ability to produce structurally coherent gene models that accurately recover protein-coding sequences. Moreover, they frequently overlook repetitive DNA sequences (repeats), despite their important roles in genome evolution and structural variation^15^, leaving the joint modeling of repeats and genes largely unexplored.

Eukaryotic gene architectures are shaped by both deeply conserved evolutionary constraints and extensive lineage-specific diversification^16–19^. Accurate gene annotation therefore requires models that can capture conserved evolutionary signals while adapting to lineage-specific variations in exon-intron organization, intron length, and repeat composition. It also requires modeling long-range genomic dependencies, as many eukaryotic genes span tens to hundreds of kilobases. However, existing deep learning models are typically restricted to local genomic contexts of approximately 10– 40 kb, limiting their ability to annotate long and structurally complex genes. Beyond these algorithmic challenges, advanced annotation models remain difficult for the broader biological community to deploy, owing to the machine-learning expertise and GPU infrastructure.

To address these limitations, we present OrionGeno, a phylogeny-aware deep-learning framework for end-to-end eukaryotic genome annotation. It integrates phylogenetic context with long-range sequence modeling under a unified prediction objective, enabling joint annotation of genes, including exons, introns, and UTRs, together with repeats directly from genomic sequence. This design removes the need for separate repeat masking or species-specific evidence integration during inference. Architecturally, OrionGeno combines bidirectional mamba blocks (BiMamba)^20^ with a U-Net-like encoder-decoder^21^, allowing the model to capture both local and long-range genomic dependencies while maintaining computational efficiency for large genomic inputs.

We demonstrate the scalability of OrionGeno by applying it to chromosome-level eukaryotic genomes from NCBI that lack reference gene annotations, generating annotations for more than 5,300 genomes and increasing public family-level annotation coverage by 47.4%. In benchmarks spanning 36 phylogenetically diverse eukaryotes, OrionGeno achieves higher accuracy than existing annotation methods across multiple biological scales, from exon recovery and gene model reconstruction to protein completeness and structural fidelity. We further show that OrionGeno-derived annotations support downstream evolutionary analyses and identify candidate protein-coding loci absent from existing protein-coding annotations even in well-curated genomes.

To broaden access and reduce technical barriers, we provide an OrionGeno ecosystem comprising a user-friendly web platform for automated annotation, programmatic API access, and a public annotation database for data exploration and reuse. Collectively, OrionGeno provides a scalable and accessible framework for eukaryotic genome annotation, helping bridge the gap between rapidly expanding genome sequencing and functional discovery across eukaryotic diversity.

## Results

### OrionGeno overview

We developed OrionGeno as a phylogeny-aware framework for *ab initio* eukaryotic genome annotation across diverse evolutionary lineages. OrionGeno requires only genomic sequences in FASTA format^22^ as input and generates standardized annotation files for gene structures, including exons, introns, and UTRs, together with repeats independent of external RNA or protein evidence (**Fig. 1a**).

**Fig 1.**
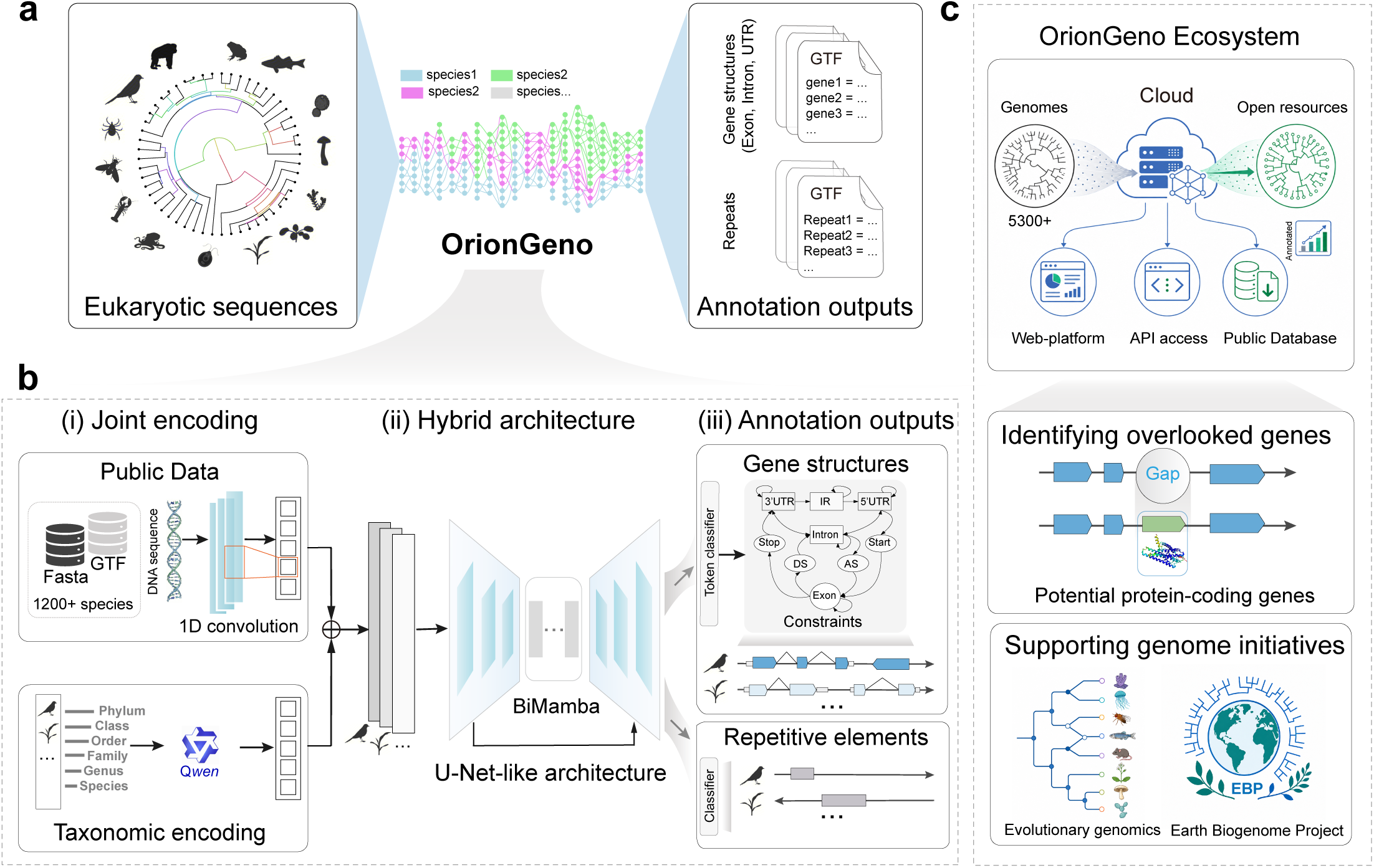
Overview of OrionGeno framework and ecosystem. **a,** Conceptual workflow. OrionGeno integrates genomic sequences (FASTA format) with hierarchical taxonomic priors, from Kingdom to Species, to generate standardized gene annotations in GTF format. The framework jointly predicts gene structures, including exons, introns, and untranslated regions (UTRs), together with repetitive elements (repeats) within a unified deep-learning architecture. **b,** OrionGeno architecture. (i) Phylogeny-augmented sequence encoding. A Qwen-based encoder converts taxonomic hierarchies into high-dimensional embeddings, which are integrated with genomic sequence features to provide evolutionary context for annotation. (ii) Multi-scale dependency modeling. A hybrid backbone combing bidirectional Mamba blocks (BiMamba) with a U-Net-like encoder-decoder captures both local sequence motifs and long-range structural dependencies. (iii) Annotation outputs. Task-specific output heads generate nucleotide-resolution probabilities for gene repeat elements. Gene predictions are further refined by a biologically constrained hidden Markov model (HMM) decoder to produce coherent gene models. **c,** OrionGeno ecosystem and applications. OrionGeno provides an integrated ecosystem consisting of a cloud-based web platform for interactive genome annotation, programmatic API access, and a public annotation database. These components enable scalable genome annotation with both user-friendly and computational interfaces. OrionGeno facilitates the identification of previously overlooked protein-coding genes and supports large-scale genome initiatives, including biodiversity genomics efforts such as the Earth BioGenome Project.

OrionGeno was trained on over 1,200 representative species (**Supplementary Table 3**), predominantly curated from the NCBI RefSeq database^23^. It employs a multi-task hybrid architecture that integrates phylogenetic context with genomic sequences to jointly predict gene structures and repeats (**Fig. 1b, Supplementary Fig. 1**). Hierarchical taxonomic metadata are encoded into phylogenetic representations (**Supplementary Fig. 2a, b**) and fused with local sequence features extracted from the input genome. This design allows the model to exploit conserved signals shared across related species while retaining information about lineage-specific variation in gene architecture and repeat composition. To capture genomic dependencies across multiple spatial scales, OrionGeno combines BiMamba^20^ with a U-Net-like encoder-decoder^21^, inspired by recent cutting-edge genomic models^24–26^. This hybrid backbone captures distal sequence dependencies while preserving fine-grained local features required for exon-intron boundary recognition. Its subquadratic computational complexity enables the context window to scale efficiently to 1 Mb, mitigating the input-length limitations of Transformer-based models.

OrionGeno was trained through a two-stage strategy that combines self-supervised pre-training and supervised fine-tuning. Inspired by the success of genomic language models (gLMs)^27–30^, we first applied a masked language modeling (MLM) objective^31^, tasking the model with reconstructing randomly masked nucleotides to learn representations of genomic sequence grammar. In the supervised stage, OrionGeno jointly optimizes gene structure annotation and repeat classification through multi-task learning. The objective combines categorical cross-entropy (CCE) with a task-aware F1-oriented loss tailored to coding regions^8^, improving learning from sparse functional elements embedded in a large non-coding background. The backbone was preserved across both stages to maintain continuity between representation learning and downstream annotation.

Beyond the core model, the OrionGeno ecosystem provides open and scalable access through an interactive web platform, programmatic APIs, an open-source repository, and a public database of OrionGeno-generated annotations (**Fig. 1c**, https://db.cngb.org/genomics/orion_geno). By integrating these components into a unified ecosystem, OrionGeno enables scalable genome annotation, lowers technical barriers, and supports broad reuse of generated gene models across the genomics community, establishing a foundation for large-scale comparative genomics and biodiversity initiatives such as EBP.

### OrionGeno enables large-scale annotation of unannotated genomes

We applied OrionGeno to unannotated chromosome-level eukaryotic genome assemblies in NCBI (**Fig. 2a**). These genomes span a broad taxonomic landscape, with major representation from classes such as Insecta, Magnoliopsida, Actinopteri, Mammalia, and Aves. OrionGeno annotated all 5,310 genomes within a week on 40 NVIDIA 5090D GPUs, demonstrating its scalability for high-throughput genome annotation.

**Fig 2.**
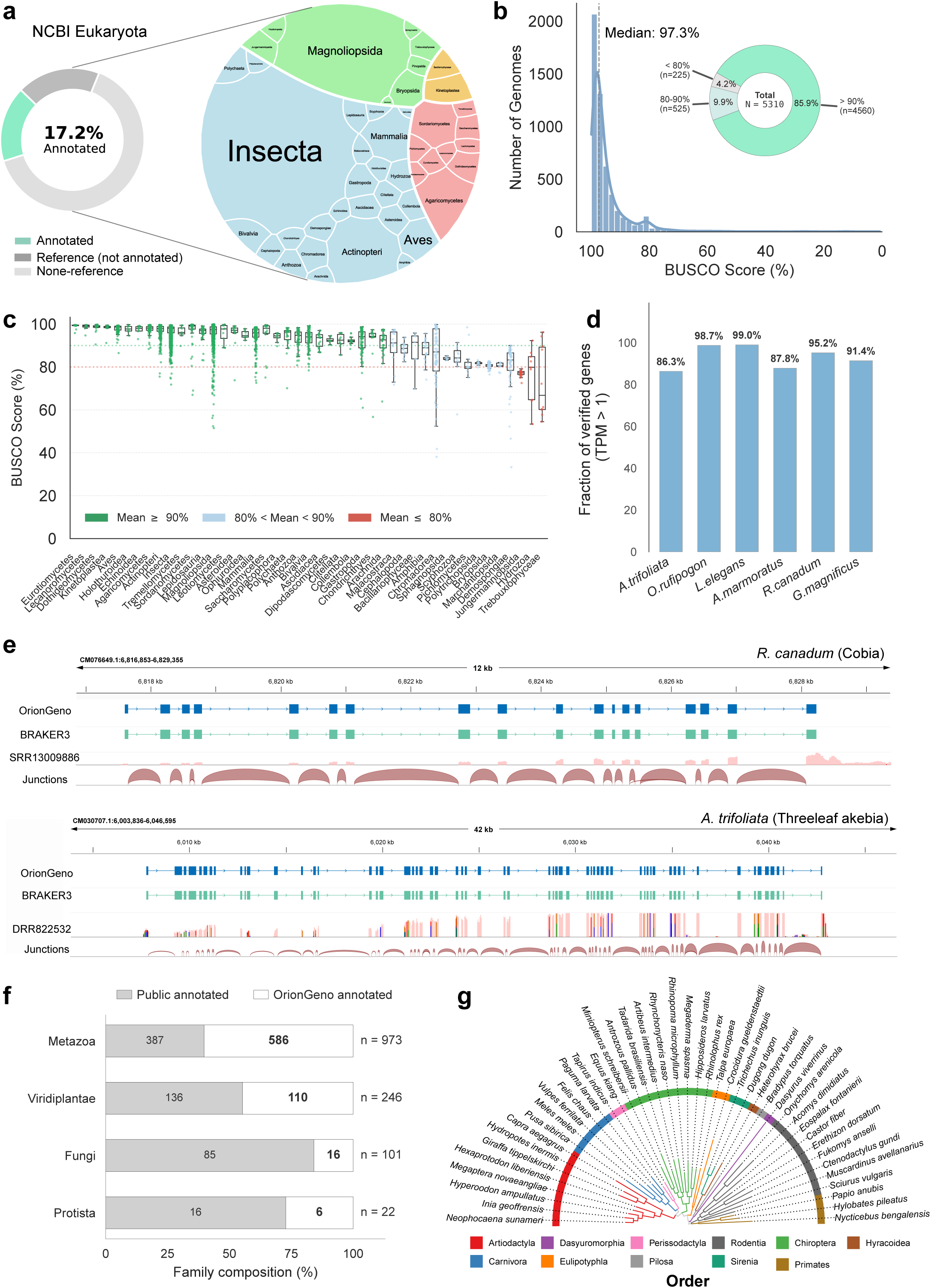
OrionGeno enables large-scale annotation of unannotated NCBI genomes. **a,** Expansion of annotated eukaryotic genome resources. Only 17.2% of NCBI eukaryotic genomes currently have functional annotations. The Voronoi treemap illustrates the taxonomic diversity of the 5,310 chromosome-level genomes lacking annotation in NCBI that were annotated by OrionGeno, with broad representation across major eukaryotic lineages, such as Insecta, Actinopteri, Aves and Magnoliopsida. **b,** BUSCO completeness. OrionGeno-predicted proteomes achieve a median BUSCO completeness of 97.3%, with 85.9% of genomes exceeding 90% completeness. **c,** Performance stability across taxonomic classes. Box plots summarize BUSCO score distributions across 45 eukaryotic classes. 30 classes (green) achieve mean BUSCO scores above 90%, whereas only 3 classes (red) fall below 80%. **d-e,** Transcriptomic validation. **(d)** RNA-seq-based validation across six representative species demonstrates high transcript support, with a mean of 93.1% of predicted genes supported by expression evidence (TPM > 1). **(e)** Representative IGV tracks in *Rachycentron canadum* and *Akebia trifoliata* demonstrate high structural concordance between OrionGeno and BRAKER3 predictions, supported by RNA-seq coverage and splice junction evidence. **f**, Expansion of family-level genome annotation coverage. OrionGeno provides annotation for 5,310 previously uncharacterized species, spanning 1,343 families. Of these, 718 families (right) currently lack any annotated representatives in public databases, highlighting the OrionGeno’s contribution to expanding genomic annotation resources. **g,** Phylogenomic application. A high-resolution phylogenetic tree reconstructed using OrionGeno-predicted gene models from representative Mammalia species illustrates its utility for phylogenomic analysis and comparative genomics.

Annotation completeness was first evaluated using Benchmarking Universal Single-Copy Orthologs (BUSCO). Across all 5,310 genomes, OrionGeno achieved a median BUSCO completeness of 97.3% (**Fig. 2b, Supplementary Table 4**). Overall, 85.9% of species (4,560 genomes) had BUSCO completeness above 90%, while only 4.2% scored below 80%. High annotation quality was broadly maintained across eukaryotic diversity. Among the 45 classes represented by more than six species, 42 had mean BUSCO scores exceeding 80%, including 30 classes with mean scores above 90% (**Fig. 2c**). High completeness was observed across vertebrates, land plants, and diverse invertebrates such as insects, arachnids, and mollusks. Lower performance was limited to several evolutionarily divergent lineages, such as Hydrozoa and Trebouxiophyceae, likely due to their substantial genomic divergence^32^, rapid evolutionary rates^33^, and the scarcity of closely related high-quality reference annotations.

To further validate the predicted gene models, we selected six representative multicellular eukaryotes, spanning vertebrates, invertebrates, and land plants, for which abundant public RNA-seq datasets are available (**Supplementary Table 5**). RNA-seq evidence supported the transcriptional activity of most predicted gene models, yielding a mean verification rate of 93.1% at TPM > 1 (**Fig. 2d**). Despite using genomic sequence along, OrionGeno rivaled BRAKER3^5^, a state-of-the-art annotation pipeline that integrates RNA and protein evidence, achieving a 4.4% relative improvement in BUSCO completeness (**Supplementary Table 6**). Representative loci from *Rachycentron canadum* and *Akebia trifoliata* showed strong structural concordance between OrionGeno and BRAKER3 annotations, together with agreement with RNA-seq coverage and splice junctions (**Fig. 2e**). Notably, OrionGeno accurately reconstructed a 60-exon gene model in *A. trifoliata* using sequence alone, highlighting its ability to recover highly complex gene models without external evidence. At scale, OrionGeno substantially expanded the phylogenetic coverage of publicly available genome annotations. Compared with existing annotation resources, including NCBI, EMBL-EBI, CNSA, and NGDC, OrionGeno generated gene annotations for 718 families with no publicly available annotated representatives, encompassing 1,387 species across diverse evolutionary lineages (**Fig. 2f**). Of these, 962 species achieved high-quality gene annotations with BUSCO completeness above 90% (**Supplementary Table 7**). By adding these newly represented families, OrionGeno increased family-level annotation coverage by 47.4%, with the largest expansions observed in Insecta (+188 families), Actinopteri (+64), Magnoliopsida (+30), Gastropoda (+26), and Bivalvia (+22) **(Supplementary Fig. 4)**. These results establish OrionGeno as a scalable framework for converting rapidly accumulating genome assemblies into high-quality and interpretable annotation resources.

### OrionGeno supports evolutionary and comparative genomics

We next evaluated the utility of OrionGeno-derived annotations for evolutionary analyses using a subset of high-quality mammalian genomes from the large-scale annotation set, selecting one representative species from each family. Using OrthoFinder^34^, we found that the majority of predicted genes were assigned to orthologous groups, with a mean clustered percentage exceeding 90% (**Supplementary Fig. 5**). OrionGeno annotations also recovered extensive sets of conserved orthologs shared across mammalian lineages, demonstrating the consistency of predicted gene models within this clade (**Supplementary Fig. 6**). Phylogenomic reconstruction based on these annotations produced a topology highly concordant with established mammalian systematics, with species correctly grouped into expected orders, including Primates, Rodentia, Chiroptera, and Artiodactyla, and other mammalian orders (**Fig. 2g**). Even for phylogenetically challenging groups, such as highly specialized aquatic mammals and the rapidly diversifying Chiroptera, the reconstructed topology maintained well resolved with strong nodal support. These results support the utility of OrionGeno-derived annotations as reusable resources for high-resolution comparative genomics.

### OrionGeno improves genome annotation accuracy across multiple biological scales

To systematically evaluate OrionGeno, we established a phylogenetically diverse benchmark of 36 high-quality genomes that were strictly excluded from model training (**Supplementary Table 8)**. Performance was assessed across multiple biological scales, from nucleotide-level classification and gene structure reconstruction to proteome completeness and protein structural fidelity (**Supplementary Fig. 3**).

Across divergent eukaryotic lineages, OrionGeno consistently outperformed existing *ab initio* gene prediction methods, including Augustus^3^, Tiberius^8^, Helixer^11^, and ANNEVO^13^, under standardized evaluation metrics^35^ (**Supplementary Table 9**). At the exon level, OrionGeno exhibited a 12.8% relative improvement in F1 score over ANNEVO, the current best-performing method at this scale in our benchmark (**Fig. 3a**). This advantage was larger at the gene level, where OrionGeno achieved a 32.0% relative increase in F1 score over the best competing method (**Fig. 3b**). OrionGeno also improved proteome completeness, with a 6.6% relative increase in BUSCO completeness and scores above 90% for nearly all the evaluated species (**Fig. 3c**).

**Fig 3.**
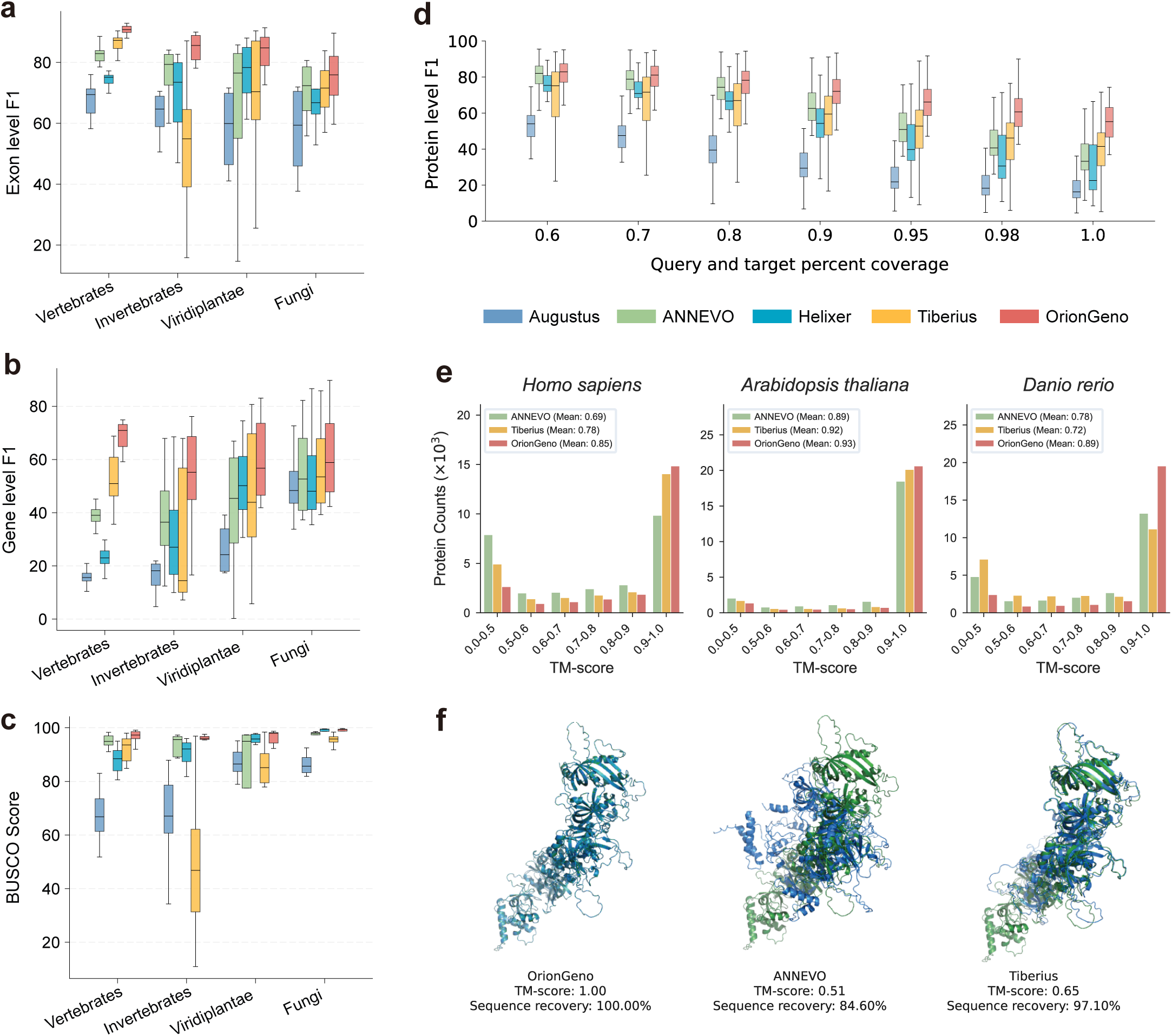
Multiscale benchmarking of OrionGeno annotation accuracy. **a-c,** Benchmarking against state-of-the-art methods. Performance across vertebrates, invertebrates, Viridiplantae, and fungi is quantified using exon-level F1 **(a)**, gene-level F1 **(b)**, and BUSCO completeness **(c)**. OrionGeno achieves improvements over state-of-the-art methods including ANNEVO, Tiberius, Helixer, and Augustus, with relative gains of 12.8%–34.8% in exon-level F1, 30.2%–180.0% in gene-level F1, and 6.6%– 32.0% in BUSCO completeness. **d,** Proteome-level reconstruction. Box plots show protein-level F1 scores under increasing alignment stringency (coverage thresholds from 0.6 to 1.0, see Methods). OrionGeno exhibits progressively increasing advantage with higher alignment stringency, reaching a relative improvement of 4.3%–37.2% across coverage thresholds from 0.6 to 1.0, where predicted proteins must fully match reference sequences. **e,** Protein structural fidelity. TM-score distributions across three model organisms show that OrionGeno yields higher structural similarity to reference proteins, with increased enrichment of predictions showing high TM scores (> 0.9). **f,** Representative structural alignments. Structural comparisons between OrionGeno predictions and benchmark methods illustrate differences in agreement with reference protein conformations. OrionGeno achieves improved structural consistency, with higher TM-scores and sequence recovery.

Given that BUSCO orthologs represent only a small fraction of the total gene set and do not account for false-positive predictions, we next evaluated protein-level accuracy across the full predicted proteome using a protein-level F1 metric (Methods). As alignment coverage requirements became more stringent, the performance gap between OrionGeno and competing methods progressively widened (**Fig. 3d**). OrionGeno showed a 4.3% relative improvement in protein-level F1 score at a coverage threshold of 0.6, and this advantage expanded to 37.2% at the strict threshold of 1.0, where predicted proteins were required to fully match reference sequences (i.e., 100% identity and 100% coverage).

We next asked whether improved gene model reconstruction translated into higher protein structural fidelity. Across three representative model organisms, OrionGeno achieved higher Template Modeling (TM) scores^36^ than competing methods, including ANNEVO and Tiberius, with mean TM-scores of 0.85, 0.93, and 0.89 in *Homo sapiens, Arabidopsis thaliana*, and *Danio rerio*, respectively (**Fig. 3e**). Consistently, OrionGeno generated more high-confidence protein models (TM-score > 0.8) and fewer low-confidence predictions (TM-score < 0.5), indicating improved recovery of structurally complete proteins. In a representative example (**Fig. 3f**), small differences in protein sequence recovery between methods resulted in substantial structural deviations among methods, whereas OrionGeno’s full-length reconstruction (1,217 aa) yielded a near-identical structural model. This case illustrates that annotation errors can be amplified at the structural level, emphasizing the importance of accurate sequence recovery for downstream functional studies. Together, these benchmarks demonstrate that OrionGeno improves genome annotation accuracy across multiple biological scales, from exon and gene reconstruction to protein completeness and structural fidelity.

### OrionGeno identifies candidate protein-coding loci absent from reference protein-coding annotations

The protein-coding catalogs of model organisms are widely regarded as the most complete owing to decades of intensive curation^23^, yet they may still remain non-exhaustive. Beyond benchmarking against known annotations, we therefore asked whether OrionGeno could identify candidate protein-coding loci absent from current reference annotations of protein-coding genes.

EggNOG-based^37^ ortholog analysis confirmed that most OrionGeno predictions were functionally concordant with RefSeq annotations. This analysis also identified 476, 198 and 242 high-confidence seed orthologs absent from reference protein-coding annotations in *Homo sapiens*, *Danio rerio*, and *Taeniopygia guttata*, respectively (**Fig. 4a**). To systematically evaluate these candidates, we focused on ortholog-supported loci with no genomic coordinate overlap with existing RefSeq or Ensembl annotations of protein-coding genes. Among these candidates, a total of 86, 64, and 52 loci, respectively, showed clear transcriptional support in public multi-tissue transcriptomic datasets^38,39^, with high expression levels (TPM > 5) (**Fig. 4b, Supplementary Tables 10–12)**. A subset of candidates, comprising three, two, and one loci in the three species, respectively, matched known protein entries in the NCBI non-redundant database^40^, providing independent support for their protein-coding potential (**Supplementary Table 13-14**). Representative loci demonstrated convergent support from dense RNA-seq coverage and consistent protein homology (**Fig. 4c**). AlphaFold-based^41^ structural modeling provided additional validation, with predicted protein forming well-ordered structures with high confidence (e.g., pLDDT > 90 for the *D. rerio* candidate) (**Supplementary Fig. 7**). Functional inspection indicated that these identified candidates appear to involve diverse biological processes, including metal-ion homeostasis and stress response in *H. sapiens* (metallothionein-like), neuronal or developmental regulation in *T. guttata* (NREP/p311-related), and receptor-mediated signaling in *D. rerio* (the GPCR GPR141).

**Fig 4.**
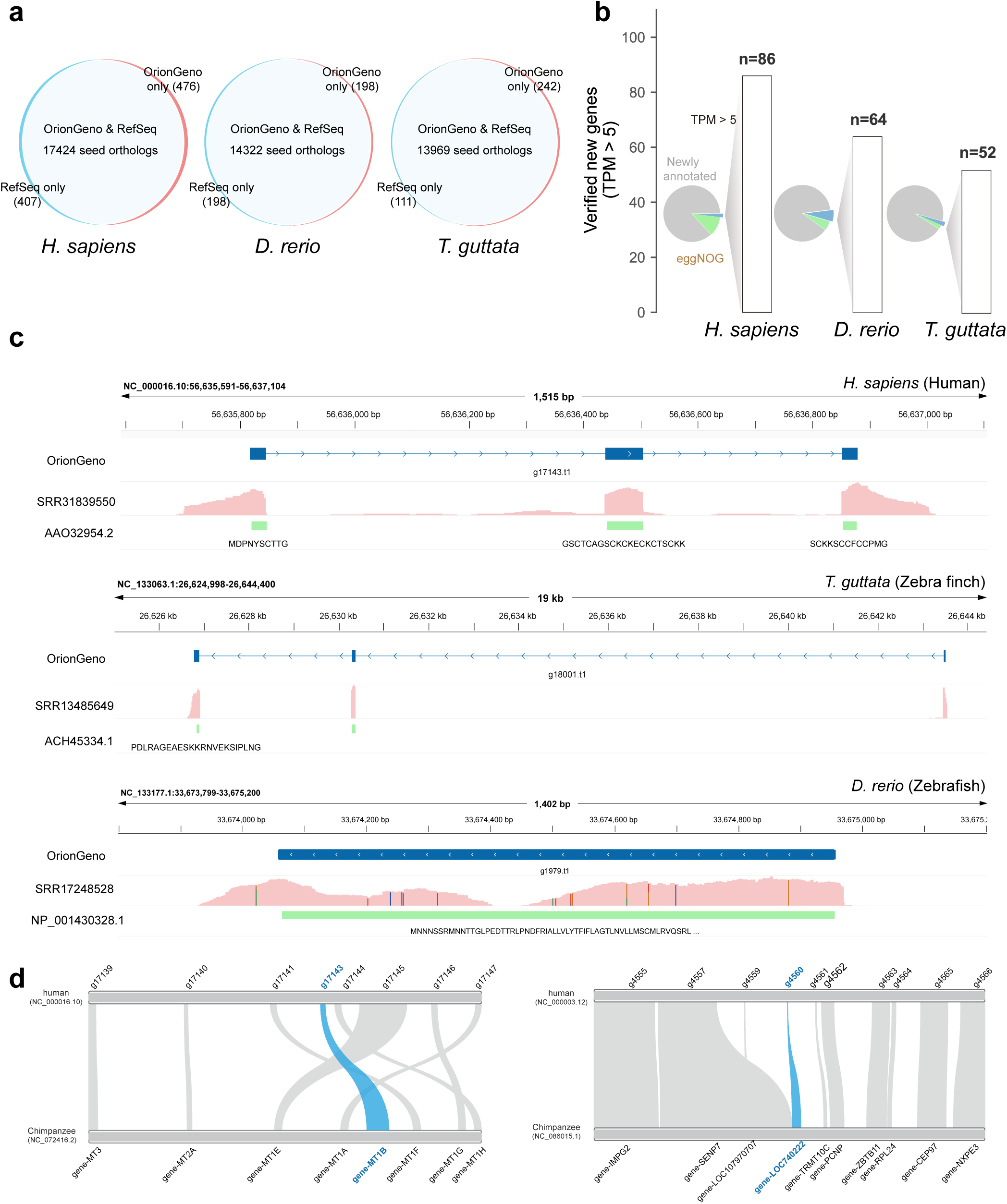
OrionGeno identifies protein-coding loci absent from reference annotations of protein-coding genes. **a,** Orthology-supported identification of loci absent from reference protein-coding annotations. Comparison of eggNOG-derived orthology assignments in *Homo sapiens*, *Danio rerio*, and *Taeniopygia guttata* shows high concordance between OrionGeno predictions with RefSeq annotations, while identifying evolutionarily conserved orthologs that are absent from current reference annotations of protein-coding genes. **b,** Transcriptomic validation of candidate loci. A stepwise filtering scheme (pie charts) classifies novel candidate loci based on their absence from RefSeq (gray), orthologous support from eggNOG (green), and transcriptional evidence (blue). Among these candidates, 86, 64 and 52 loci, respectively, show transcriptional support (TPM > 5) and no genomic coordinate overlap with existing RefSeq or Ensembl protein-coding gene annotations. **c,** Representative OrionGeno-identified protein-coding loci. IGV tracks display representative newly identified protein-coding loci in *H. sapiens*, *D. rerio*, and *T. guttata*. These predicted gene models are supported by RNA-seq evidence and homologous protein sequence matches. **d,** Syntenic conservation of OrionGeno-identified human loci with chimpanzee protein-coding genes. Blue highlights indicate OrionGeno-identified human loci and their corresponding chimpanzee protein-coding counterparts (*gene-MT1B* and *gene-LOC740222*).

We further compared the three human candidate loci supported by both transcriptional and protein-level evidence with their orthologous regions in the chimpanzee genome. The corresponding chimpanzee loci were annotated as protein-coding genes in RefSeq, whereas the orthologous human regions lack protein-coding annotations. The OrionGeno-predicted human loci were embedded within conserved syntenic blocks and maintained collinearity with neighboring orthologous genes (**Fig. 4d**). This conserved syntenic context, together with the transcriptional and protein-level evidence, supports their protein-coding potential and suggests that they may represent previously unrecognized or potentially misclassified protein-coding loci in humans.

### OrionGeno captures sequence rules underlying gene architecture

To move beyond black-box prediction, we employed an *in silico* mutagenesis strategy to elucidate the sequence determinants that drive OrionGeno annotation outputs (Methods). Across genomic positions, perturbations within exons exerted a significantly greater impact on model predictions than comparable mutations in intronic or intergenic regions. Feature contribution maps revealed that the strongest model responses were concentrated within exons, with pronounced enrichment at exon boundaries (**Supplementary Fig. 8**). Start and stop codons were among the most influential sites, producing sharp and localized signals in the contribution maps.

Targeted mutagenesis further showed that OrionGeno responds to canonical coding signals in a biologically coherent manner. For instance, mutating the initial adenine of the canonical start codon ATG of human gene *GAPDH* to alternative nucleotides (T, C, or G) triggered a reconfiguration of the predicted gene model (**Fig. 5a**). In response to the loss of this primary initiation signal, OrionGeno skipped the first two exons, classifying them as intergenic regions until a downstream ATG was encountered to reinitiate translation. By contrast, substituting the canonical stop codon TAA with the alternative stop codon TAG had little effect on the predicted gene model (**Fig. 5b**), indicating that the model recognizes the functional equivalence of multiple termination signals. Mutating TAA to a non-stop triplet, such as TAT or TAC, abolished proper translation termination and extended the predicted gene model downstream until an in-frame termination signal (TAA) was encountered. These results indicate that OrionGeno has learned sequence principles that reflect core constraints of protein-coding gene architecture.

**Fig 5.**
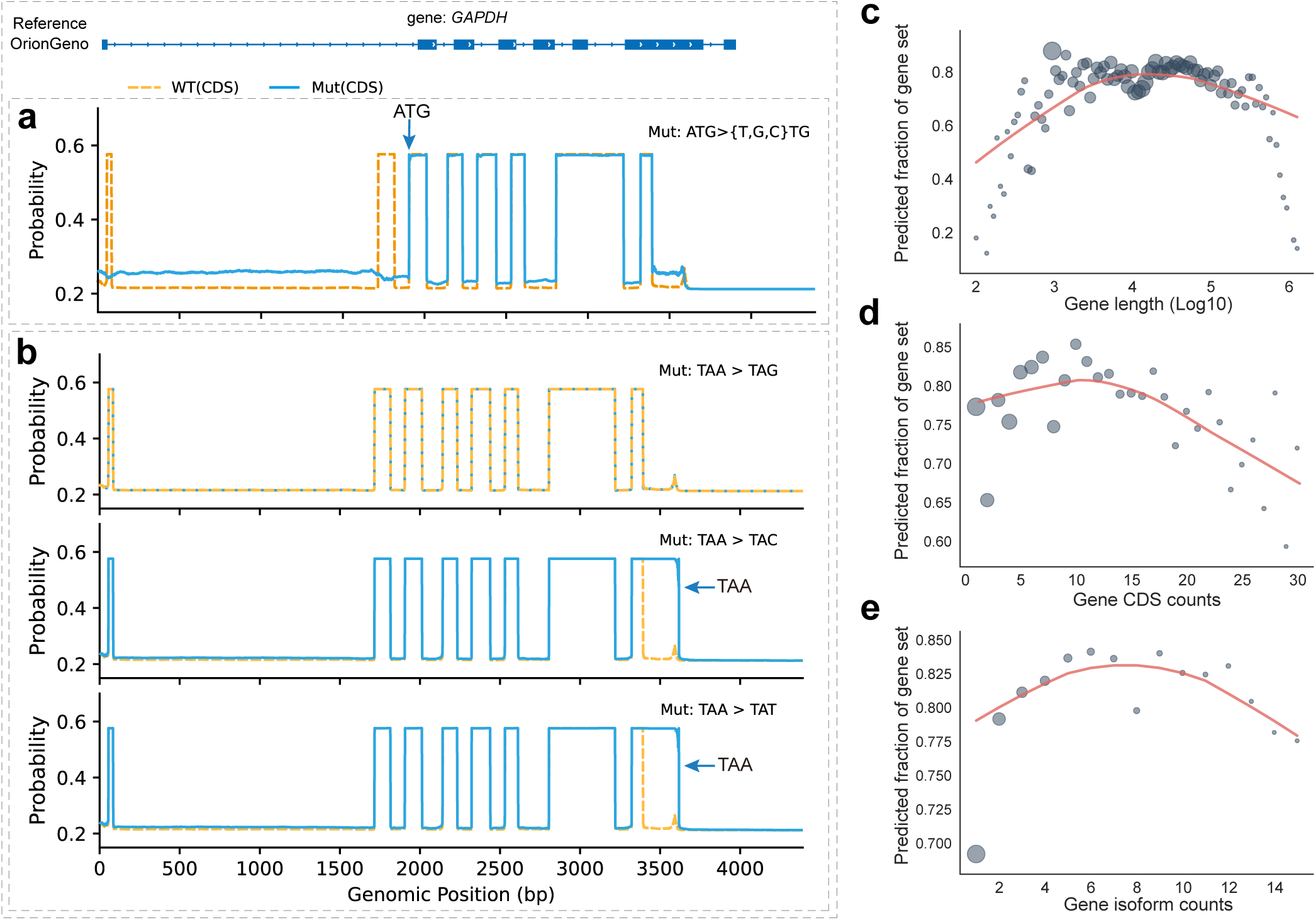
Mechanistic interpretability and robustness of OrionGeno. **a-b,** Sequence determinants underlying OrionGeno predictions revealed by in silico mutagenesis. Probability tracks illustrate model responses to targeted perturbations in the human *GAPDH* locus. Mutation of the canonical start codon (ATG→{TTG, CTG, GTG}) disrupts recognition of the initial coding region and causes downstream recovery of the gene model only after a downstream initiation signal is encountered **(a)**. In contrast, synonymous stop-codon substitution (TAA→TAG) preserves the predicted coding structure, whereas stop-loss mutations (TAA→TAC/TAT) extend the predicted coding region until the next in-frame termination signal **(b)**. These perturbation experiments demonstrate that OrionGeno captures biologically constrained sequence rules governing gene structure prediction. **c-e,** Robustness across diverse gene architectures. Scatter plots show the fraction of correctly recovered gene models across human genes stratified by genomic length **(c)**, CDS count **(d)**, and transcript isoform diversity **(e)**.

### OrionGeno is robust across diverse gene architectures

We next evaluated OrionGeno across the structural diversity of the human protein-coding genome. We stratified genes by genomic span, CDS segment count, and isoform diversity, and observed robust performance across most of the protein-coding repertoire (**Fig. 5c-e**). In particular, OrionGeno was broadly stable for loci spanning approximately 3 kb to 316 kb (10^3.5^–10^5.5^ bp), with reduced recovery observed primarily for genes at the short and long extremes of the length distribution (**Fig. 5c**). Performance was similarly stable for genes containing up to approximately 15 CDS segments and for loci associated with up to 10 annotated transcript isoforms (**Fig. 5d, e**). Beyond these ranges, the fraction of predicted genes declined progressively, most notably for genes containing more than 20 CDS segments and for loci with the greatest isoform diversity. This trend is consistent with the increasing structural complexity of these loci, for which accurate gene-model reconstruction requires the coordinated resolution of larger numbers of splice junctions and alternative transcript configurations. Nevertheless, OrionGeno maintained strong performance across the majority of human protein-coding gene architectures, supporting its robustness for large-scale genome annotation.

### OrionGeno resolves structurally challenging gene loci

Lastly, we examined OrionGeno at representative loci that capture three sources of difficulty in eukaryotic gene annotation: ultra-long genomic span, dense gene clustering, and extensive isoform diversity (**Fig. 6**). These loci require annotation models to integrate local splice and coding signals while maintaining coherent gene-level structure over long genomic distances. At the human *CNTNAP5* locus, which spans approximately 895 kb and contains 23 coding exons, OrionGeno reconstructed a continuous gene model across the full locus. By contrast, the benchmarked methods recovered only partial structures or split the locus into incomplete models, highlighting the difficulty of linking widely separated exons across an extended genomic interval.

**Fig 6.**
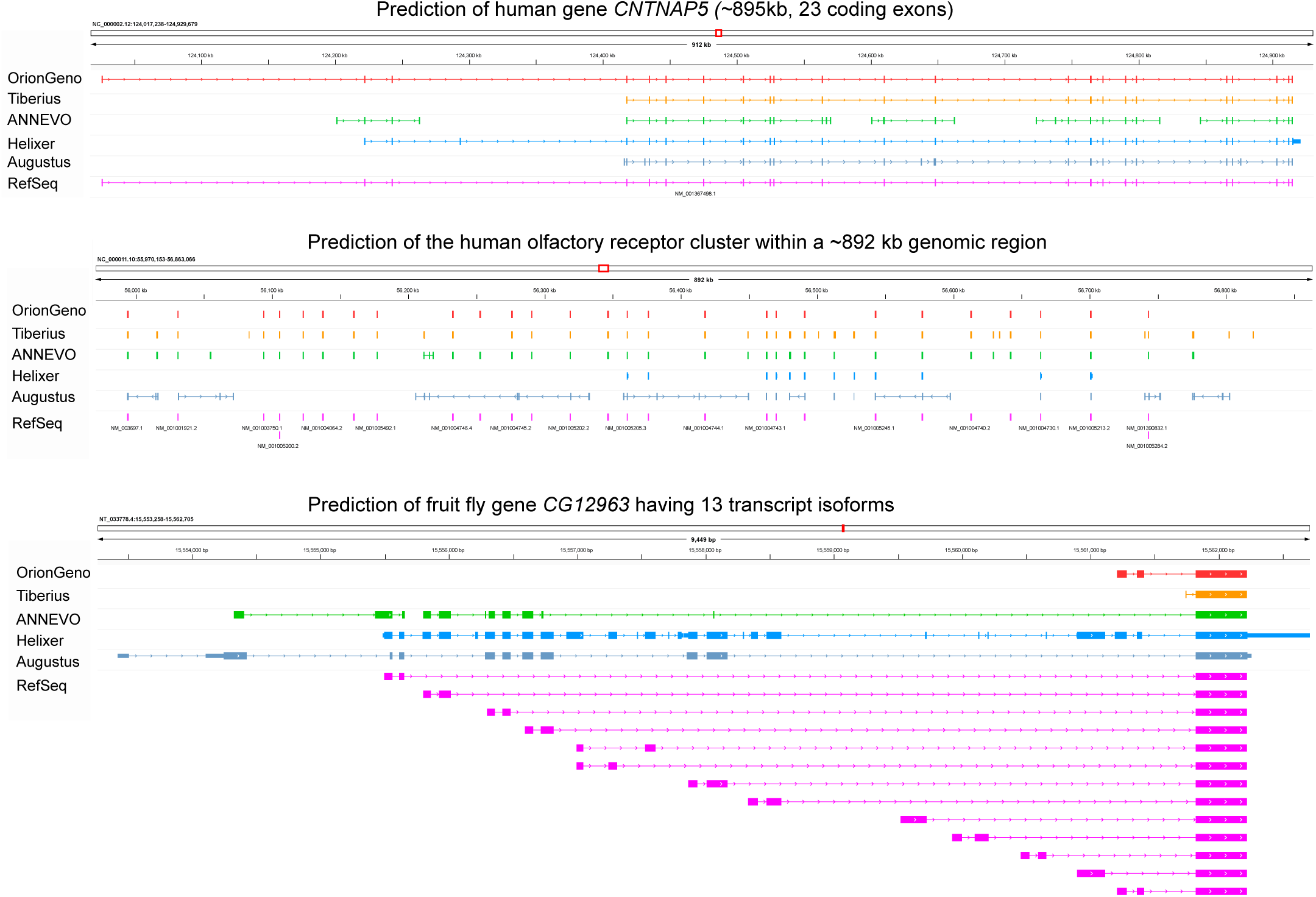
OrionGeno resolves challenging gene architectures. Representative IGV tracks illustrate loci with exceptionally long genomic spans, dense tandem gene clustering and extensive transcript isoform diversity. **Top**, the human *CNTNAP5* locus (∼895 kb, 23 coding exons), for which OrionGeno reconstructs a continuous gene model across the full locus, whereas benchmark methods produce fragmented or incomplete predictions. **Middle**, a representative ∼892-kb region of the human olfactory receptor cluster, in which OrionGeno separates individual gene units within a dense paralogous cluster. **Bottom**, the *Drosophila melanogaster* gene *CG12963*, which contains 13 annotated transcript isoforms, where OrionGeno recovers the coding structure of the longest RefSeq isoform. These examples illustrate OrionGeno’s ability to preserve long-range gene structure and resolve complex gene boundaries across diverse genomic contexts.

Paralogue-rich gene clusters present a distinct annotation challenge, particularly when closely spaced genes are encoded by single coding exons and therefore provide limited splice-structure cues for defining gene boundaries. In a representative 892-kb human olfactory receptor cluster, part of the largest and most complex tandemly duplicated gene family in the human genome^42^, OrionGeno accurately separated adjacent gene units and produced boundaries that closely matched the RefSeq annotation. By contrast, competing methods frequently missed individual genes or produced fragmented and merged models, indicating reduced robustness in resolving densely packed paralogous loci.

Isoform-rich loci present an additional source of ambiguity because multiple alternative exon configurations must be reconciled into a coherent coding model. For the *Drosophila melanogaster* gene *CG12963*, which has 13 annotated transcript isoforms, OrionGeno recovered the coding structure corresponding to the longest RefSeq isoform and precisely matched the reference annotation. By contrast, benchmarked tools were frequently misled by splicing complexity, resulting in truncated or misassembled gene models.

Together, these examples show that OrionGeno can resolve gene structures across genomic contexts that challenge both local signal detection and long-range structural consistency. This capability benefits from its hybrid architecture, in which U-Net-like convolutions capture local sequence features whereas BiMamba blocks integrate information across extended genomic contexts. Training on sequence windows ranging from 50 kb to 1 Mb further exposes the model to the spatial scales required to resolve complex eukaryotic gene architectures.

## Discussion

This study presents OrionGeno as a scalable framework for annotating eukaryotic genomes across broad phylogenetic diversity. By applying OrionGeno to more than 5,300 chromosome-level NCBI genomes without exiting gene annotations, we generated a public annotation resource and demonstrated that end-to-end genome annotation can be performed consistently at a scale that is difficult to achieve with evidence-dependent pipelines. Together with a web platform, programmatic APIs, and an integrated annotation database, OrionGeno provides reusable infrastructure for converting newly assembled genomes into functional resources, with particular value for data-poor lineages that contribute substantially to the annotation gap affecting nearly half of NCBI reference genomes^43^.

A key feature of OrionGeno is its multispecies, phylogeny-aware modeling strategy, which makes species diversity informative rather than merely heterogeneous. Eukaryotic genomes share core coding and splicing principles but differ substantially in exon-intron organization, intron density and length, splice-site context, and sequence composition^44^. By incorporating taxonomic representations during sequence modeling, OrionGeno can learn conserved signals across related species while keeping lineage-specific annotation patterns distinguishable. This design allows increased training diversity to broaden the information available to the model, supporting more stable annotation across diverse eukaryotic groups.

An important finding of this study is that OrionGeno improves annotation accuracy not only at the nucleotide and gene levels but also at the protein sequence and structural levels. This multiscale improvement is important because errors in exon connectivity can propagate into fragmented transcripts, incomplete coding sequences, and incorrectly predicted proteins. Although approximate recovery of gene-containing regions may be sufficient for broad evolutionary surveys, many downstream biological and translational applications, including functional genomics, genome editing and molecular breeding, require accurate coding models because protein domains and structural integrity can directly influence target selection and experimental design. OrionGeno’s gains at the gene, protein and structural levels indicate that improved gene model reconstruction can translate into more complete protein recovery and greater structural fidelity, thereby supporting more reliable downstream biological interpretation.

Current end-to-end gene prediction models often focus on gene structures alone, despite the importance of repeats in genome organization, evolution, and annotation error^45,46^. By jointly modeling gene structures and repeats, OrionGeno eliminates the need for separate repeat masking and improves discrimination between coding and repeat-derived signals. Consistent with this, predicted repeats showed only minimal (0.64%) overlap with consensus coding regions in the human genome, suggesting that the model can limit false-positive gene predictions in repeat-rich regions (**Supplementary Fig. 14**).

Despite these advances, several limitations remain. Reduced accuracy in basal metazoans such as *Hydra vulgaris* highlights the challenge of modeling genomes across extreme evolutionary distances, where the scarcity of closely related genomic representatives constrains model adaptation. UTRs also remain more difficult to predict than coding regions, likely owing to their lower evolutionary conservation and heterogeneous quality of UTR annotation in reference datasets^47^. In addition, the current framework reports only the longest protein-coding transcript per locus and does not yet model the full complexity of alternative splicing. Future extensions that incorporate comparative genomic signals, multi-omics evidence, and isoform-level modeling should help capture the broader functional complexity of eukaryotic genomes and expand the scope of predictive genome annotation.

Going forward, we envision OrionGeno as a scalable engine for eukaryotic genome interpretation that can operate independently of any wet experimental datasets. Ongoing efforts will extend this resource to the growing number of unannotated assemblies in NCBI and support large-scale initiatives such as EBP^2^. By combining scalable inference with public data release and accessible annotation tools, OrionGeno may help turn rapidly accumulating genome assemblies into more usable biological resources.

## Methods

### Data collection and processing

#### Data download

A total of 2,058 high-quality eukaryotic genome assemblies and their corresponding annotations were retrieved from the NCBI Reference Sequence (RefSeq) FTP site (https://ftp.ncbi.nlm.nih.gov/refseq/release/). Data from 42 additional Viridiplantae species, including bryophytes and algae, were incorporated from NCBI (https://ftp.ncbi.nlm.nih.gov/genomes/all/GCA/) and CNCB (https://ngdc.cncb.ac.cn/genbase). The NCBI Genome database was systematically screened to identify chromosome-level or complete genome assemblies without official gene annotations (December 2025), yielding a curated set of 5,310 assemblies for downstream application of OrionGeno. Scientific names and their complete taxonomic hierarchy (Kingdom to Species) for 22,444 eukaryotic species were obtained from the NCBI Genome and Taxonomy databases (https://ftp.ncbi.nih.gov/pub/taxonomy/)^48^. RefSeq accessions and taxonomy information are detailed in the **Supplementary Tables 1-2**.

#### Data usage

RefSeq species were grouped into major phylogenetic clades: vertebrates (Mammalia, Aves, Actinopterygii, Amphibia, Reptilia, and others), invertebrates (Arthropoda, Arachnida, Insecta, Cnidaria, Lophotrochozoa, and others), Viridiplantae (Monocots, Eudicots, Gymnosperms, Bryophytes, Chlorophyta), and Fungi. Species were selected to prioritize high-quality genomes (BUSCO > 90%) while maintaining balanced representation across these clades. For lineages with limited high-quality genome resources, species with lower BUSCO completeness were also included to ensure evolutionary diversity, such as *Symsagittifera. roscoffensis* (GCF_963678635.1) and *Convolutriloba. macropyga* (GCF_964194025.1). In total, OrionGeno was trained and validated on genome assemblies from 1,282 species, including 393 vertebrates, 185 invertebrates, 99 Viridiplantae, 556 fungi, and 49 protists (**Supplementary Table 3**). The test dataset comprised 36 well-curated representative species spanning major phylogenetic clades, including *Homo sapiens*, *Arabidopsis thaliana*, *Danio rerio*, *Danaus plexippus* and *Caenorhabditis elegans* etc. (**Supplementary Table 8**). All data from these test species were strictly excluded from training and validation to prevent data leakage and ensure rigorous benchmarking.

#### Data processing

High-confidence transcripts were curated by applying a series of filtering criteria. Transcripts were excluded if they met any of the following conditions: (1) presence of internal stop codons; (2) coding sequence lengths not divisible by three (i.e., non-triplet length); (3) coding sequences shorter than 60 nucleotides; or (4) overlaps with neighboring genes on the same strand. For each gene, transcript with the longest coding sequence was selected to generate unambiguous training labels. This labeling strategy aligns with OrionGeno’s single-label-per-base design, which does not model alternative splicing isoforms. For repeat prediction, training labels were derived from RefSeq soft-masked genome sequences, in which lowercase bases denote repetitive regions and uppercase bases denote non-repetitive regions. Genome sequences were segmented into chunks ranging from 51,200 bp to 1 Mb.

### Model architecture

#### Taxonomy-augmented sequence embedder

Genomic sequences were encoded using a one-hot representation with a five-token nucleotide vocabulary (*A*, *T*, *G*, *C*, *N*). An input sequence of length *L* was represented as a binary matrix *X* ∈ {0,1}^5×*L*^, which was subsequently projected into a high-dimensional feature space (*d=384*) via a one-dimensional convolutional layer.

Phylogenetic context was incorporated using taxonomic embeddings derived from the large language model Qwen-7B (https://huggingface.co/Qwen/Qwen-7B). For each species in the compiled NCBI eukaryotic taxonomy dataset, the full taxonomic hierarchy—Kingdom, Phylum, Class, Order, Family, Genus, and Species—was formatted as a structured text to generate species-specific latent representations. This procedure produced a high-dimensional embedding matrix of size (22444 × 4096), capturing hierarchical and phylogeny-aware relationships across eukaryotes. To align these taxonomic features with sequence representations, Principal Component Analysis (PCA) was applied to reduce the embedding dimensionality to *d=384*. UMAP visualization of these resulting embeddings revealed distinct clusters consistent with known evolutionary lineages, indicating that Qwen effectively captured the underlying phylogenetic structure in eukaryotes (**Supplementary Fig. 2**). The resulting taxonomic vectors were then integrated with sequence-derived features and propagated through a U-Net–inspired hybrid architecture. The taxonomy-augmented representation enables OrionGeno to reconcile gene structure variability across divergent evolutionary lineages, producing species-specific predictions tailored to the target genome.

#### Hybrid modeling backbone

OrionGeno employs an efficient hybrid architecture that integrates U-Net-like convolutional encoder-decoder^21^ with bi-directional Mamba blocks (BiMamba)^20^, supporting input sequences of up to 1 Mb in length. Convolutional layers are designed to capture local sequence dependencies, while BiMamba blocks model long-range relationships with subquadratic computational scaling. The architecture comprises three core components: (1) a U-Net-like convolutional encoder, (2) BiMamba blocks, (3) a convolutional decoder with skip (residual) connections from the encoder.

**(1) Down-sampling convolutional encoder.** The encoder consists of three successive down-sampling convolutional blocks, each followed by max-pooling with stride 2, progressively reducing the resolution of taxonomy-augmented sequence features while expanding the receptive field. This hierarchical design captures sequence patterns across multiple genomic scales, from nucleotide-level motifs to longe-range contextual dependencies. Intermediate feature maps are extracted prior to each pooling step, corresponding to effective resolutions of 2 bp, 4 bp, and 8 bp, and are retained as skip connections for the U-Net-like decoder to facilitate accurate boundary localization during reconstruction. For example, down-sampling compresses a 51,200-bp input sequence into a latent representation of length 3,200, corresponding to one token per 16 bp, before passing it to the downstream BiMamba blocks.
**(2) BiMamba.** Following the encoder, OrionGeno adopts BiMamba^20^ as its core sequence modeling module. Unlike transformer-based hybrid architectures such as Enformer^24^, Borzoi^49^, and AlphaGenome^26^, which are limited by the quadratic computational burden of self-attention, the Mamba framework enables efficient modeling of long sequences with subquadratic scaling. BiMamba extends Mamba^50^ with bidirectional processing and reverse-complement (RC) equivariance, allowing effective modeling of long-range dependencies in genomic sequences. This design achieves a favorable balance between computational efficiency and representational capacity for large-scale genomic inputs.
**(3) Up-sampling convolutional decoder.** Following the standard U-Net architecture^21^, a convolutional decoder mirroring the encoder is used to progressively restore base-pair resolution from the 16 bp contextual embeddings. The decoder consists of three up-sampling convolutional stages, with skip connections that fuse upsampled features with corresponding encoder representations at each scale. This design integrates long-range contextual information with high-resolution local features, enabling accurate reconstruction of genomic structures. Feature channels are gradually reduced across layers, yielding a final 384–channel representation at single-base-pair resolution for downstream base-wise prediction.

#### Gene and repeat output heads

At the final stage of OrionGeno, the single-base-resolution embeddings generated by the decoder are processed by two independent point-wise linear classification heads. This dual-head design enables multi-task learning, allowing the model to simultaneously predict gene structural elements and repeat regions. The gene annotation head projects the per-base representations into an eight-class logit vector, followed by Softmax normalization to produce posterior probabilities for exon, intron, splice donor site, splice acceptor site, start codon, stop codon, 3′ UTR, and 5′ UTR. The repeat head performs binary classification at each nucleotide position, predicting whether each site belongs to repetitive or non-repetitive genomic regions. The two heads are jointly optimized during training.

#### Gene structure decoder

To decode canonical exon–intron architectures and enforce reading-frame consistency, OrionGeno incorporates a hidden Markov model—a classical paradigm in gene finding—that constrains state transitions to biologically plausible paths, inspired by Helixer^11^ and Tiberius^8^. The postprocessor enforces canonical DNA patterns at exon boundaries by constraining state-specific emissions: the START state exclusively emits the ATG codon; the STOP state emits TAG, TAA, or TGA with equal probability (0.33 each); donor sites emit GT (0.99) or GC (0.01); and acceptor sites deterministically emit AG (1.0). During inference, the optimal state sequence is decoded using the Viterbi algorithm^51^ with fixed, untrained HMM parameters, and the resulting gene structures are exported as GTF annotations. The model contains approximately 121 million trainable parameters, with ∼11% in the encoder, ∼74% in the BiMamba blocks, and ∼15% in the decoder. The HMM post-processing module is not trained, and its parameters remain fixed during inference. All the codes were implemented in PyTorch (version 2.7 with CUDA 12.6).

### OrionGeno training

OrionGeno was trained using a two-stage learning strategy comprising (1) self-supervised pretraining with a BERT-style masked language modeling (MLM) objective to capture general sequence patterns across diverse eukaryotic genomes and (2) supervised fine-tuning for gene and repeat prediction. The model architecture remained unchanged across stages, facilitating the transfer of learned sequence representations to downstream functional tasks.

#### Pretraining with masked nucleotides modeling

Pretraining followed a MLM paradigm, in which the model was tasked with reconstructing randomly masked nucleotides within a genomic sequence. 20% of nucleotide positions in each input sequence were dynamically selected, among which 50% were replaced with a

<MASK> token, 25% were substituted with random nucleotides, and the remaining 25% were left unchanged. The model was trained to predict the original nucleotides at the selected positions using a token-level cross-entropy loss ℒ_MLM_(*θ*):

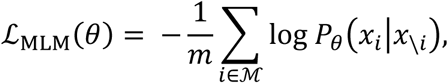

where *θ* denotes the model parameters, ℳ is the set of selected nucleotide positions, and *m* = ⌈ℳ⌉. *x_i_* represents the true nucleotide at the *i*-th masked position, and *P_θ_*(*x_i_*|*x*_\*i*_) indicates the probability assigned by the model to the true nucleotide *x_i_* given its contextual sequence (*x*_\*i*_).

#### Supervised training for gene prediction

Supervised fine-tuning was employed to learn base-wise genomic annotations from labeled datasets. To address the extreme sparsity of exonic regions, we employed a tailored CCE-F1-loss^8^, which combines categorical cross-entropy (CCE) with an F1-based for sequences containing exon labels or false positive rate (FPR) objective term for sequences without exon labels. Specifically, the F1-loss ℒ_F1_ is formulated as:

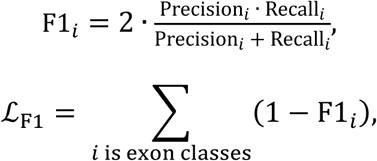

where F1*_i_* denotes the estimated F1-score for class *i*, computed as the harmonic mean of the estimated Precision*_i_* and Recall*_i_*. In sequences lacking exon labels, the F1-loss cannot be computed. In such cases, an estimated false positive rate (FP̂R) is used:

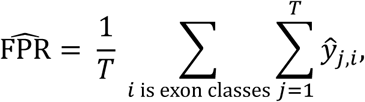

where *T* is the sequence length and *ŷ_j,i_* is the predicted probability of class *i* at position *j*. The total loss ℒ_CCE−F1_ is then defined as:

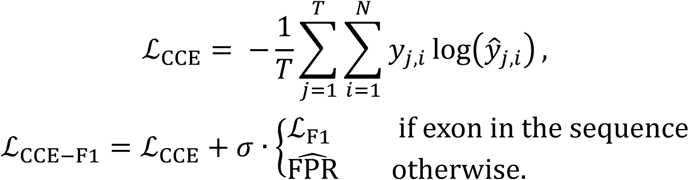

Here, *N* denotes the total number of classes, *y_j_*_,*i*_ is the true label for class *i* at position *j*, and *ŷ_j,i_* is the corresponding predicted probability. A weighting factor *σ* balances the contribution of F1-based (or FPR) objective relative to the CCE term and was set to 2 in our model.

#### Supervised training for repeat prediction

Supervised fine-tuning for repeat prediction was formulated as a binary classification task at single–base-pair resolution. For each nucleotide position, the model predicts the probability of belonging to repeat or non-repeat regions. The repeat prediction head was optimized using a binary cross-entropy (BCE) loss:

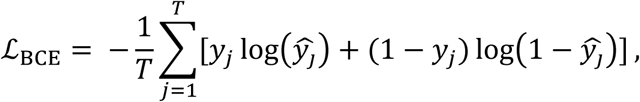

where *T* denotes the sequence length, *y_j_* ∈ {0, 1} is the ground-truth label at position *j*, and *ŷ_j_* is the predicted probability of being a repeat.

### Benchmarking against state-of-the-art methods

OrionGeno was benchmarked against state-of-the-art *ab initio* gene prediction methods, including Augustus^3^, Tiberius^8^, Helixer^11^, and ANNEVO^13^. Among these, Augustus represents a classical HMM-based approach, whereas the others are deep learning-based methods. To ensure a fair comparison, test species were primarily selected from the original benchmark datasets used in the Tiberius, ANNEVO, and Helixer studies. For clades not covered in their benchmarks, additional high-quality model organisms were included, see all the test species in **Supplementary Table. 4.**

For each test species, whole-genome annotations were generated using all evaluated methods. For each tool, we preferentially used models trained on the same species or on closely related species (**Supplementary Fig. 11**), following the recommended usage of each method to ensure comparability across approaches. SegmentNT^12^ was excluded from the large-scale benchmark because it produces base-level multi-label predictions rather than discrete gene structures, making it incompatible with standard GTF-based evaluation. Instead, we performed qualitative visualization on a selected set of human genes, for which gene structures were manually reconstructed from SegmentNT outputs (**Supplementary Fig. 12**).

#### Benchmark metrics

Gene annotation accuracy was evaluated across nucleotide, exon, gene, protein, and protein structure levels. Nucleotide-level performance was assessed using genome-wide F1 scores, providing a base-pair-resolution measure of labeling precision. To evaluate gene structure reconstruction across the exon and gene levels, predictions were benchmarked against RefSeq annotations using standardized evaluation metrics^35^, specifically utilizing gffcompare^52^ to compute exact-match precision, recall, and F1 scores for exon and gene structures. To account for alternative splicing, a predicted transcript was considered as a true positive if it achieved a perfect structural match with any annotated isoform at the corresponding locus.

At the protein level, proteome completeness was first evaluated using BUSCO scores^53^. Given that BUSCO is restricted to highly conserved core orthologs and does not penalize false-positive predictions, we additionally implemented a comprehensive protein-level F1 metric, adapted from ANNEVO^13^, to evaluate the full predicted gene repertoire while explicitly accounting for false positives. A predicted gene model was categorized as a True Positive (TP) at the protein level only when it simultaneously satisfied the following three stringent criteria: (1) protein sequence homology, where the best-matching segment between the predicted and reference protein sequences met predefined DIAMOND^54^ thresholds (bit-score ≥ 50 and e-value ≤ 1e-5), retaining only the top-scoring hit per protein; (2) genomic coordinate consistency, requiring overlap between predicted and reference genes at the same locus; (3) alignment coverage, where both query coverage (fraction of the predicted protein aligned) and target coverage (fraction of the reference protein aligned) exceeded a predefined threshold (60%–100% in this study). Based on counts of true positives (TP), reference positives (P), and predicted positives (PP), protein-level F1 score is defined as follows:

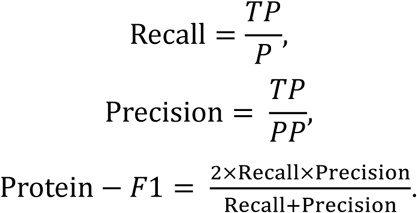

At the protein structural level, we evaluated the structural fidelity of predicted proteomes using Template Modeling scores (TM-scores)^36^. Protein sequences derived from RefSeq annotations and from annotations generated by OrionGeno, Tiberius, and ANNEVO were independently folded using ESMFold^55^ under identical configurations. Structural similarity between predicted and reference models was quantified using TM-score, with higher values indicating greater agreement in global tertiary structure.

### *In silico* mutagenesis analysis

To investigate how OrionGeno decodes genomic elements, we performed an *in silico* mutagenesis analysis inspired by previous studies^56,57^. This analysis quantifies how nucleotide substitutions at one genomic position affect the model’s prediction at another position. Specifically, for a given input sequence, we mutated the nucleotide at a query position *i* to each of the three alternative bases and measured the resulting change in the predicted probability at a target position *j*, expressed as a log-odds ratio. For a sequence of length *N*, the contribution score *s _i_*_,*j*_ between query position *i* and target position *j* is defined as:

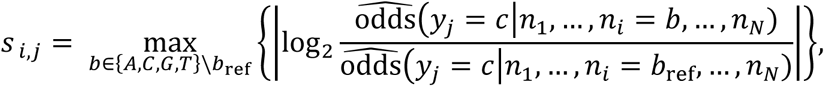

where *b*_ref_ denotes the reference nucleotide at position *i*, *y_j_* is the predicted label at position *j* and *c* is the label of interest. The term ôdds denotes the odds estimated from the model’s predicted probabilities. Contribution scores are computed for all position pairs (*i*, *j*) with *i* ≠ *j*, thereby excluding self-dependencies.

This procedure can be applied to all query–target position pairs and provides a systematic measure of the extent to which predictions at a target position depend on perturbations at a query position while all other nucleotides are held constant.

## Data availability

Genomic assemblies and annotations used for model training, benchmarking, and applications are listed in Supplementary Tables 1-3, and are available from RefSeq (https://ftp.ncbi.nlm.nih.gov/refseq/release/), NCBI (https://ftp.ncbi.nlm.nih.gov/genomes/all/GCA/), and CNCB (https://ngdc.cncb.ac.cn/genbase). Taxonomic hierarchies are available at https://ftp.ncbi.nih.gov/pub/taxonomy/. RNA-seq accessions are provided in Supplementary Table 5 and accessible via the ENA (https://www.ebi.ac.uk/ena/browser/home). All *ab initio* gene annotations generated by OrionGeno have been deposited in CNGB under project ID CNP0009228 (https://db.cngb.org/data_resources/project/CNP0009228).

## Code availability

The OrionGeno model can be accessed via GitHub at https://github.com/BGIResearch/OrionGeno.

## Acknowledgements

This work is supported by the National Key R&D Program of China (grant no. 2024YFA0919700), the National Natural Science Foundation of China (grant nos. 32500577 to L.L. and 32370666 to Q.Y.L.). We acknowledge the State Key Laboratory of Genome and Multi-omics Technologies, Stomics Cloud platform (https://cloud.stomics.tech/), and Sugon for providing computational resources. We thank S.S.D., Y.L.Z, and G.Y.F. for initial discussion and support.

## Author contributions

P.Y. and L.L. conceptualized the study. P.Y., S.F.W., L.L., X.D.C., and C.Z. were responsible for model design and implementation. Y.W.W., J.Y.W., and K.S. performed data collection and preprocessing. X.D.C. performed the benchmark and downstream analysis. Y.D., Y.L.P., X.P.Z., and H.P.X. contributed to the RNA and protein evidence. L.L. and X.D.C. were responsible for figure visualization and typesetting. L.L. wrote the original manuscript, and all authors contributed to refinement. T.W., Y.L. provided scientific insights and technical advice. P.Y., Y.X.L., Y.Z., Q.Y.L., and X.X. co-supervised the study.

