## Supplementary Figures 1-14 for "Scaling genome annotation across the eukaryotic tree of life with OrionGeno"

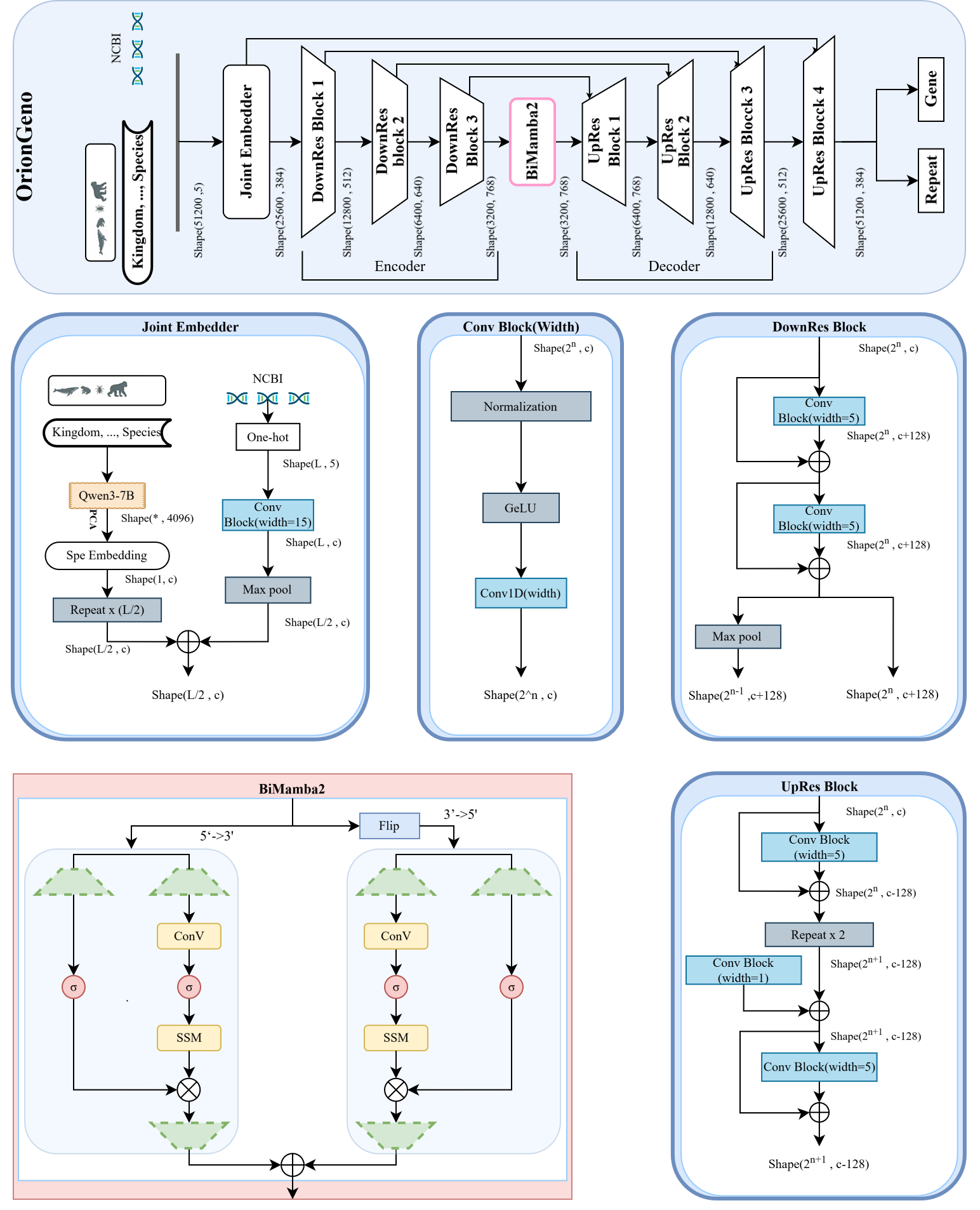

**Supplementary Fig. 1 | OrionGeno model architecture.** **Top,** overall model architecture. OrionGeno integrates genomic sequence features with species-level phylogenetic context through a joint embedding module, followed by a hybrid encoder–decoder backbone composed of residual downsampling blocks (DownRes), a bidirectional Mamba module (BiMamba), and residual upsampling blocks (UpRes). The model generates two position-wise outputs corresponding to gene annotation and repeat prediction. **Joint Embedder,** Species information, represented as a hierarchical taxonomic string from kingdom to species, is encoded by Qwen3-7B and projected into a species embedding, which is broadcast along the sequence axis and fused with genomic sequence features. In parallel, genomic DNA is one-hot encoded and processed by a convolutional stem followed by max pooling. The two branches are combined to produce phylogeny-aware sequence representations. **Conv Block and DownRes Block,** The basic convolutional block consists of normalization, GeLU activation and 1D convolution. DownRes blocks apply stacked convolutional operations with residual connections, followed by max pooling, to progressively expand channel capacity while reducing sequence length in the encoder. **BiMamba2,** The central BiMamba2 module models long-range genomic dependencies in both the 5′→3′ and 3′→5′ directions. Sequence features are processed in parallel by directional state-space branches and then merged, enabling bidirectional context modeling while preserving computational efficiency. **UpRes Block.** In the decoder, UpRes blocks progressively restore sequence resolution through repeated upsampling and convolutional refinement, while integrating skip connections from the encoder to recover fine-scale positional information. Together, these modules enable OrionGeno to jointly resolve gene structures and repetitive regions from raw genomic sequence and phylogenetic input.

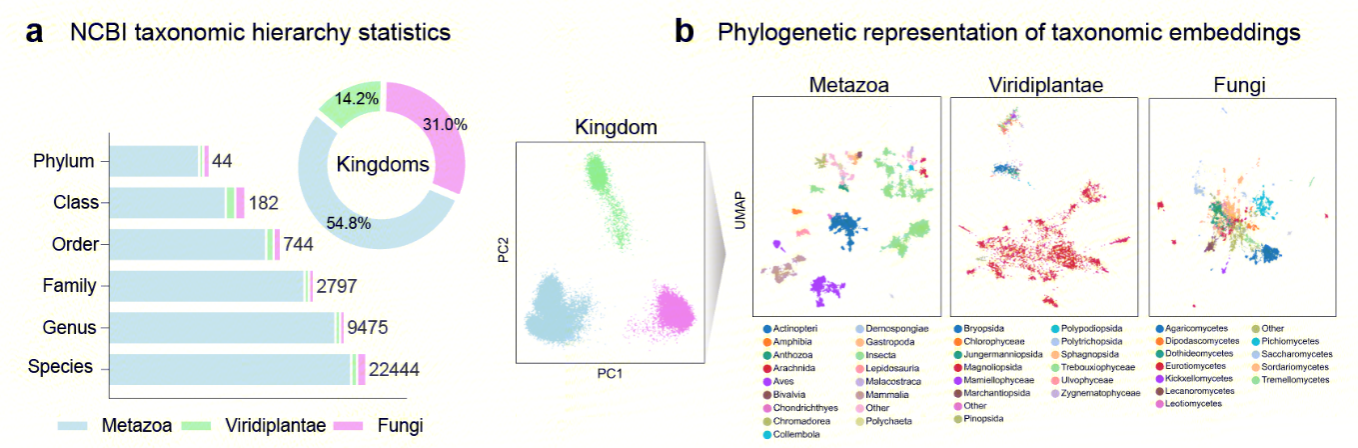

**Supplementary Fig. 2 | Taxonomic coverage and visualization of the Qwen-derived taxonomy embeddings. a,** Taxonomic hierarchy statistics and kingdom-level embedding structure. The number of unique taxa represented at each rank in the NCBI taxonomy dataset, spanning phylum, class, order, family, genus and species across Metazoa, Viridiplantae and Fungi. Donut chart showing the kingdom-level composition of the species set. **b,** Phylogenetic representation of taxonomic embeddings. Principal component analysis (PCA) of the learned taxonomic embeddings at the kingdom level, showing clear separation among the three major eukaryotic groups. Uniform Manifold Approximation and Projection (UMAP) visualization of the taxonomic embeddings within Metazoa, Viridiplantae and Fungi. Points are colored by lower-level taxonomic groups, showing that the embeddings preserve phylogenetic structure and organize species into coherent lineage-resolved clusters.

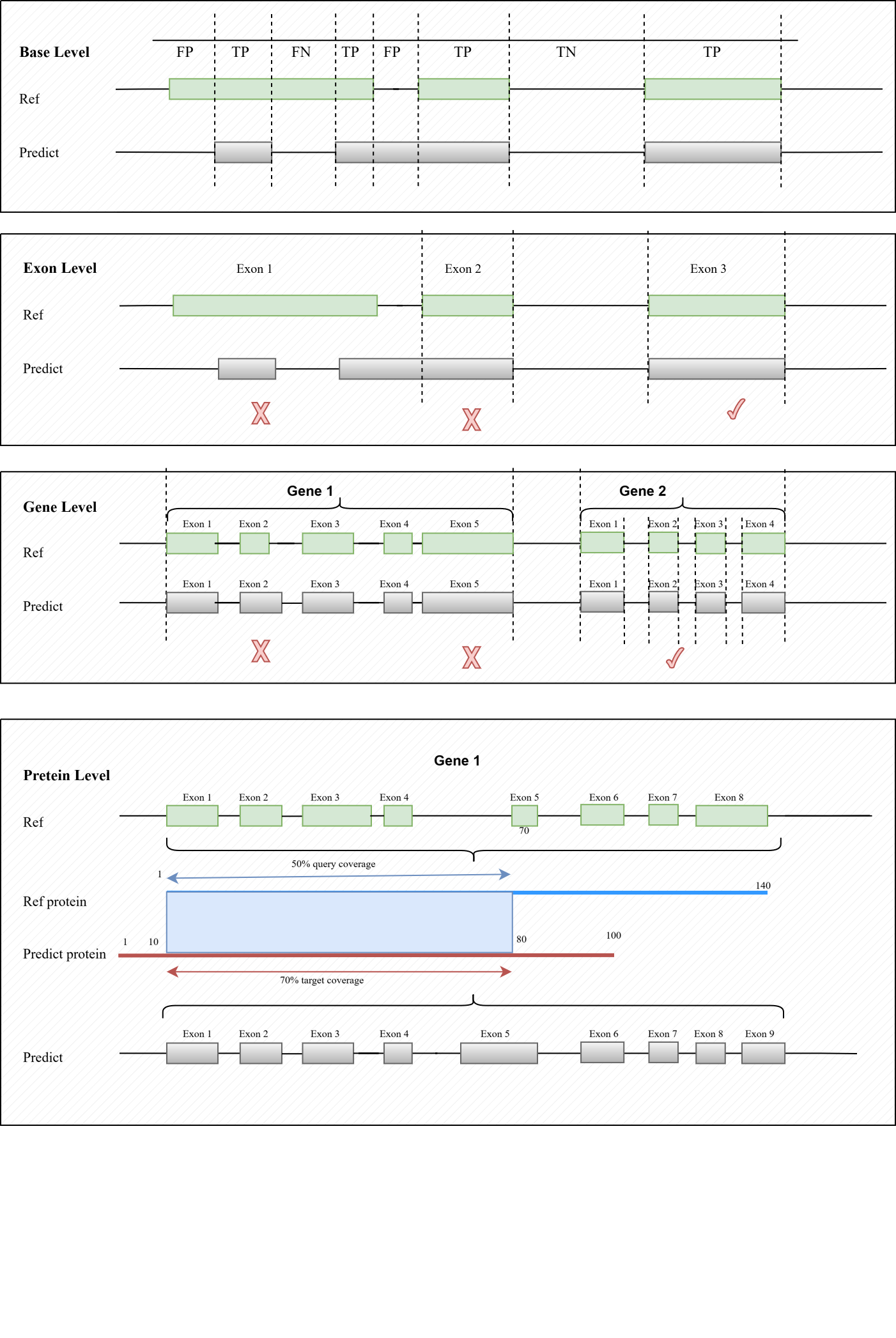

**Supplementary Fig. 3 | OrionGeno model architecture.** Multi-level benchmarking framework for evaluating gene annotation quality. Schematic illustration of performance assessment at four complementary levels. **Base level,** predictions are evaluated at single-nucleotide resolution by comparing base-wise labels to the reference, yielding TP, FP, FN and TN assignments. **Exon level,** individual exons are evaluated using exact-match criteria, such that only predicted exons with perfectly matched coordinates are counted as correct. **Gene level,** complete exon–intron structures are compared, and a predicted transcript or gene is considered correct only if its full structure exactly matches the reference. **Protein level,** predicted proteins are matched to reference proteins based on sequence homology, genomic overlap and bidirectional alignment coverage, allowing assessment of coding-sequence recovery beyond exact structural concordance.

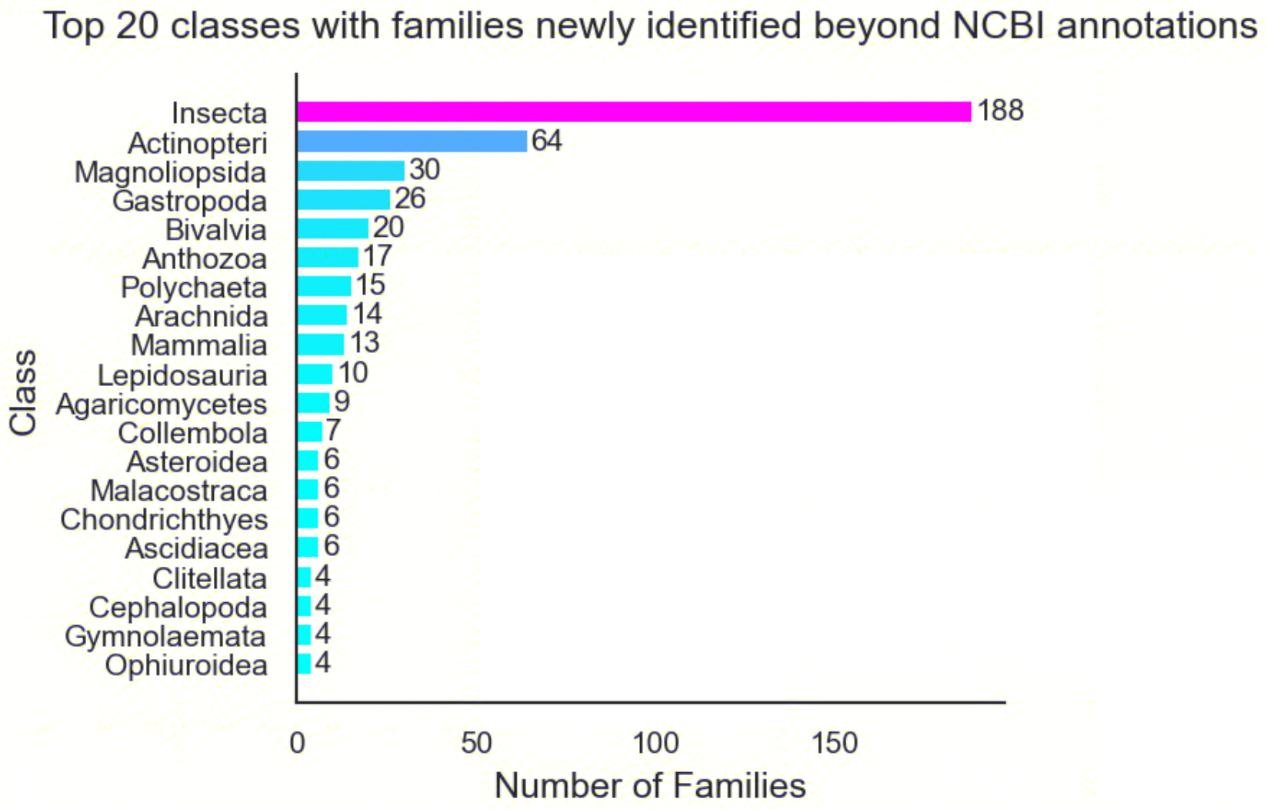

**Supplementary Fig. 4 | Structural characteristics and taxonomic expansion of high-confidence OrionGeno annotations. a,** Distributions of gene, intron and CDS lengths across major eukaryotic phyla annotated by OrionGeno, shown on a log10 scale (BUSCO > 90%). **b,** Numbers of families for which OrionGeno achieved high-confidence annotations (BUSCO >90%) in Metazoa, Viridiplantae and Fungi. **c,** Top 20 classes contributing newly annotated families beyond current NCBI coverage, illustrating the broad phylogenetic expansion enabled by OrionGeno.

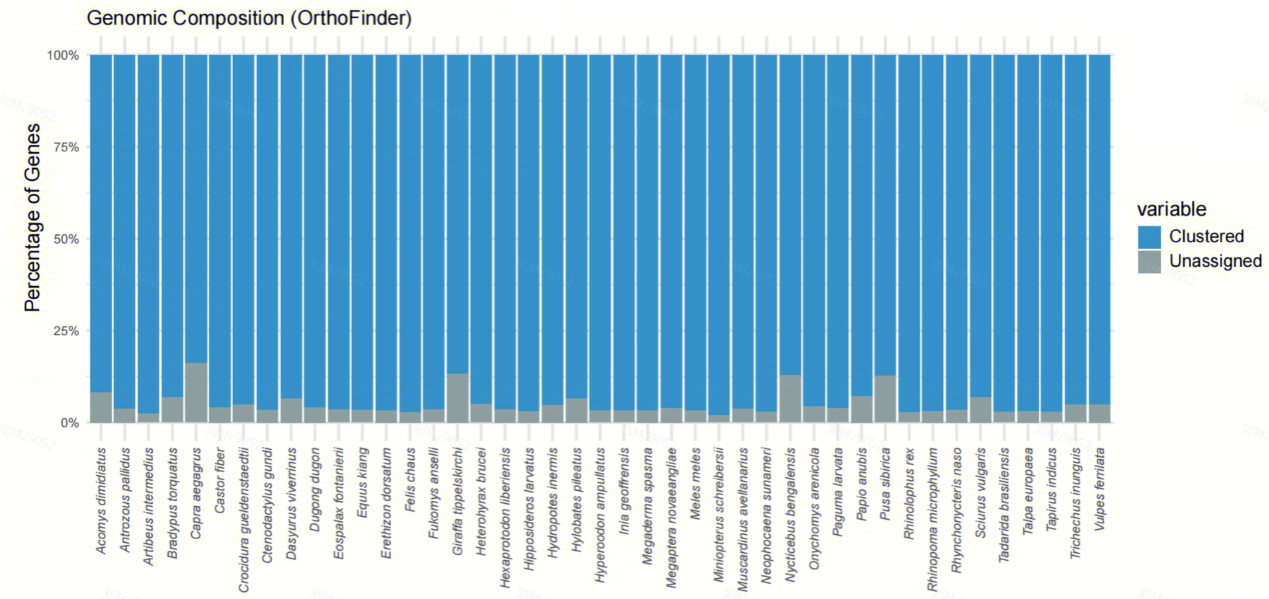

**Supplementary Fig. 5 | Genomic composition defined by OrthoFinder orthogroup assignment.** For each species, stacked bars show the percentage of predicted genes assigned to orthogroups (Clustered) or left unassigned by OrthoFinder. Most genes were incorporated into orthogroups across species, supporting the suitability of OrionGeno annotations for comparative genomics and phylogenomic analysis.

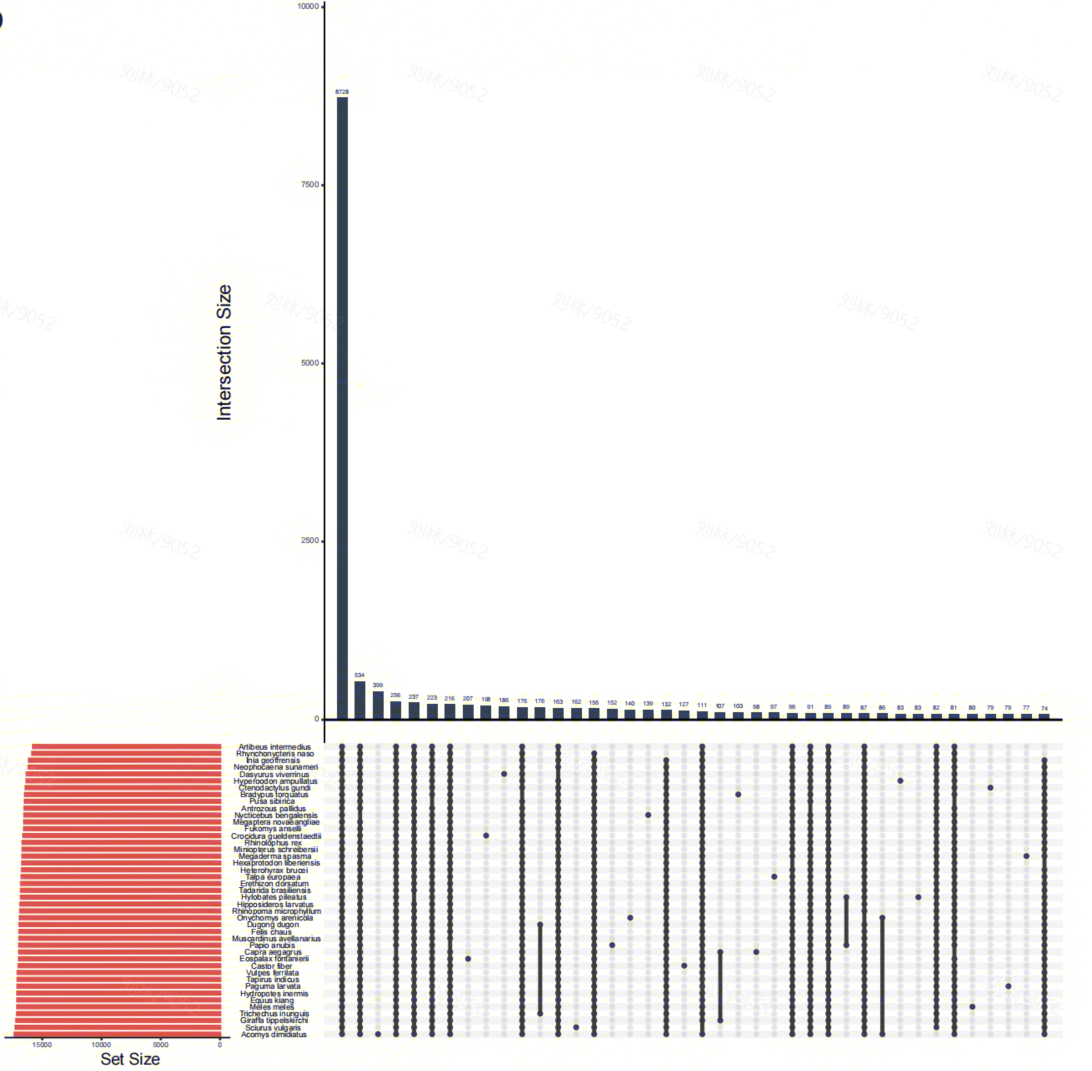

**Supplementary Fig. 6 | UpSet visualization of orthogroup overlap across species.**

Horizontal bars show the total number of orthogroups detected in each species. Connected dots indicate the species composition of each intersection, and vertical bars show the corresponding intersection sizes.

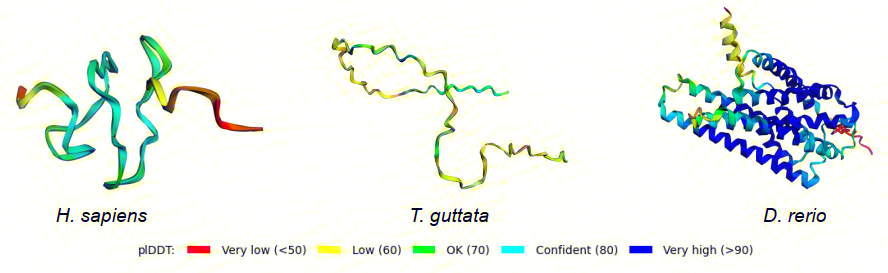

**Supplementary Fig. 7 | Structural support for coding potential.** Predicted three-dimensional structures of the novel gene products shown in (Figure. 3c), colored by pLDDT score. The prevalence of highly ordered folding (pLDDT > 90, blue), particularly the globular architecture in *D. rerio*, supports their protein-coding potential.

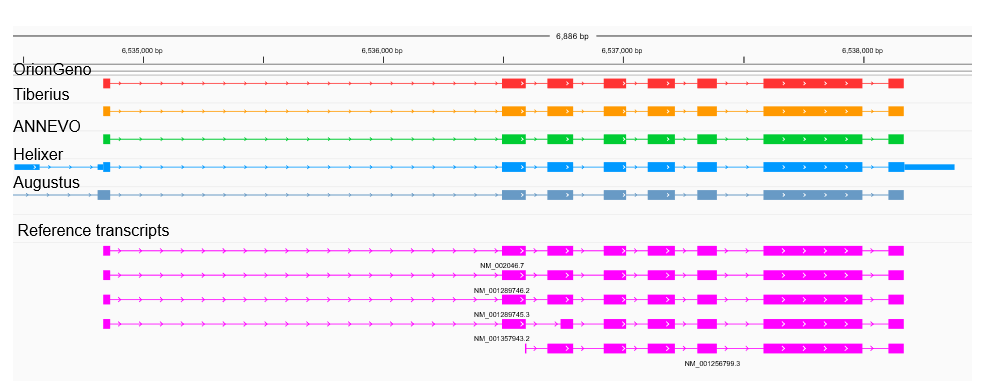

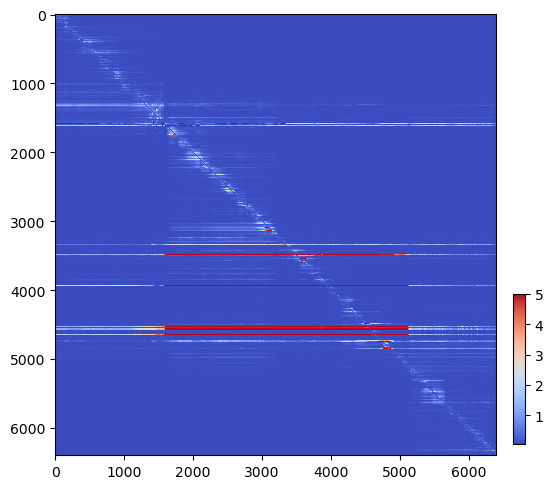

**Supplementary Fig. 8 | Structural accuracy and dependency mapping at a representative locus. a,** Comparison of OrionGeno and existing methods at a representative alternatively spliced locus. OrionGeno shows closer agreement with the reference transcript structures than Tiberius, ANNEVO, Helixer and AUGUSTUS.
**b,** *In silico* mutagenesis heatmap for the same region, showing how single-nucleotide perturbations at one position alter annotation predictions at other positions. Strong off-diagonal signals reveal long-range dependencies underlying exon connectivity and transcript structure prediction.

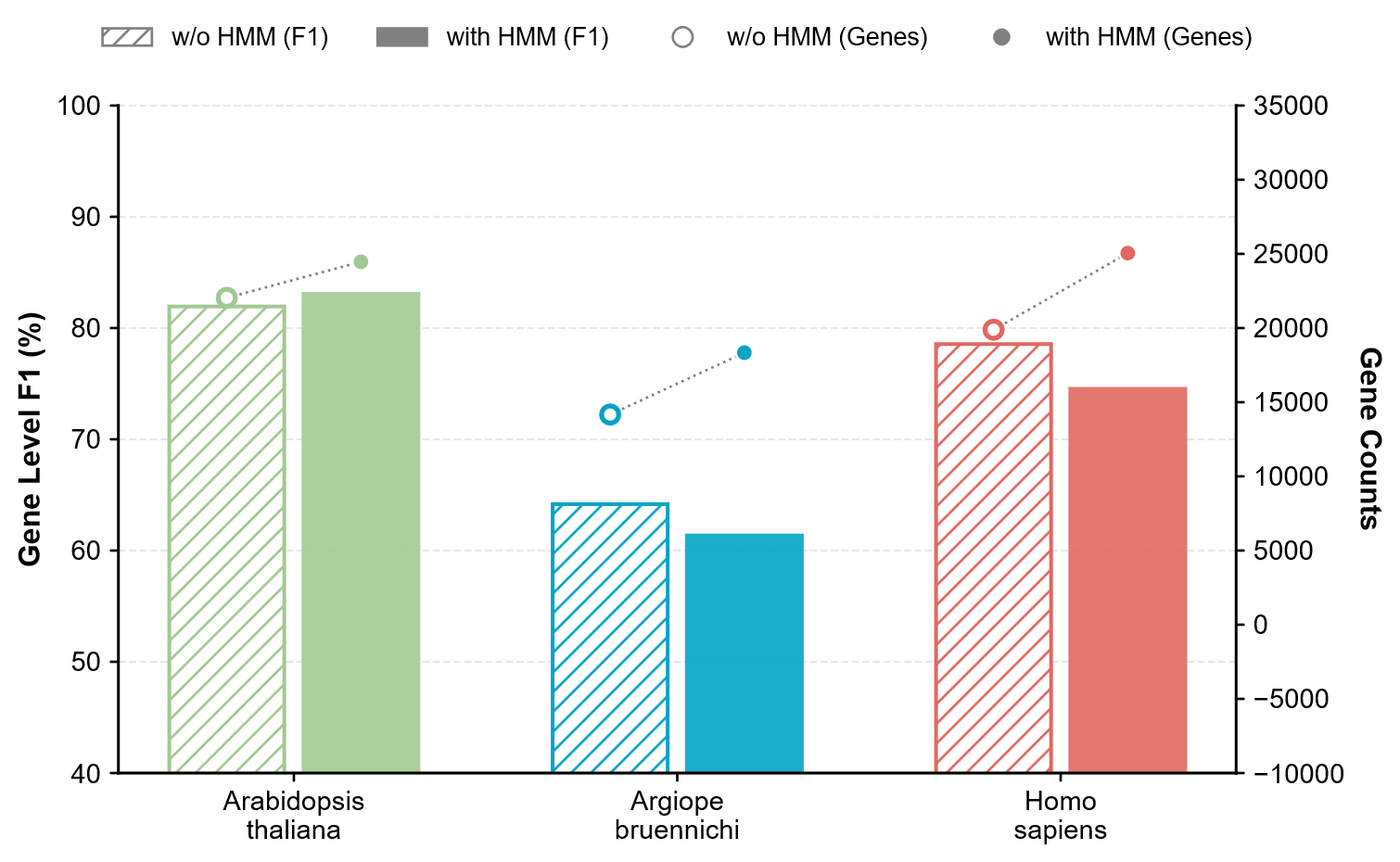

**Supplementary Fig. 9 | Effect of HMM-based decoding.** Gene-level F1 scores (bars, left y axis) and numbers of predicted genes (circles, right y axis) for *Arabidopsis thaliana*, *Argiope bruennichi* and *Homo sapiens*, comparing predictions with and without the HMM decoder. Hatched/open symbols denote predictions without HMM, and solid/filled symbols denote predictions with HMM.

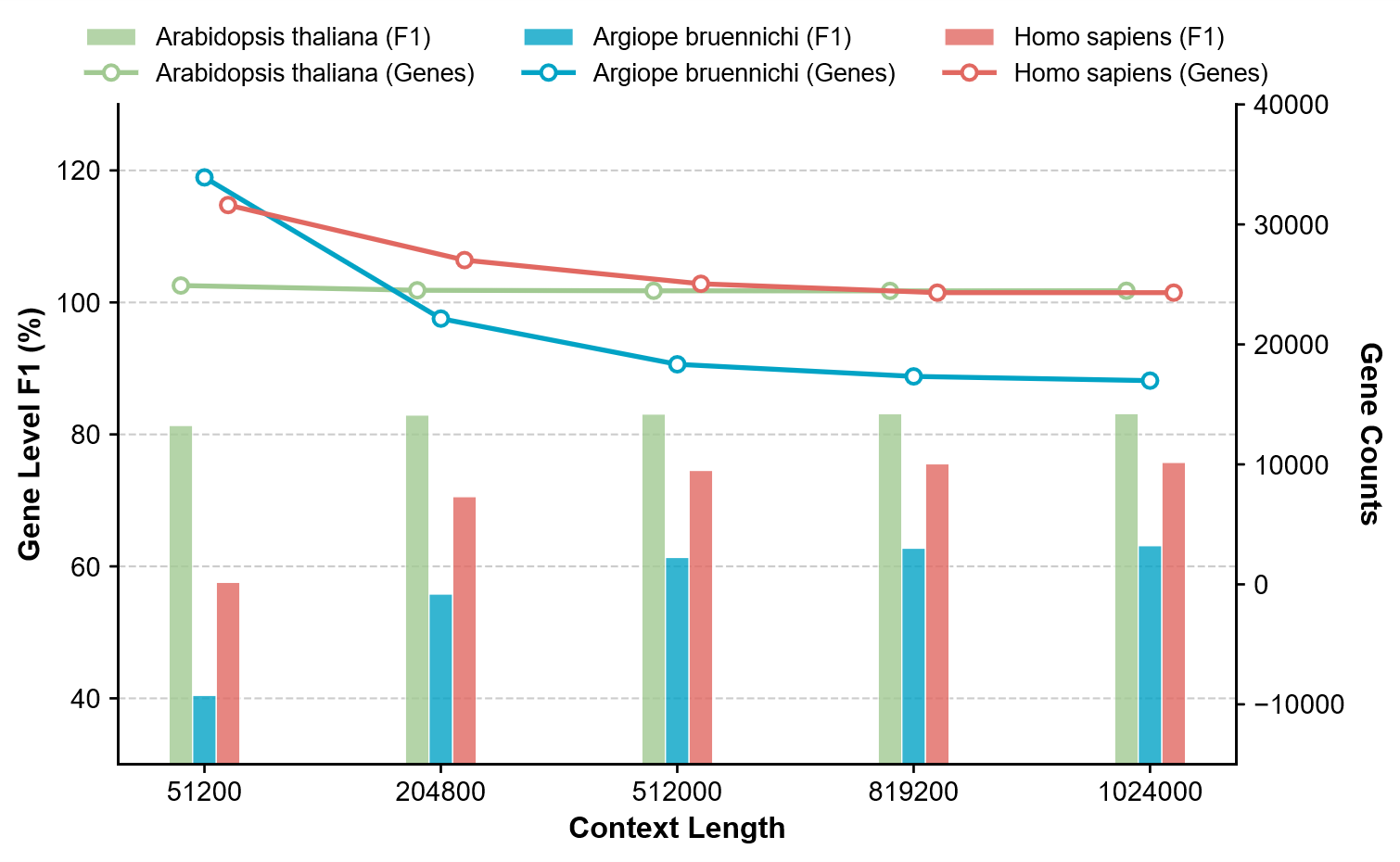

**Supplementary Fig. 10 |** **Effect of inference context length on annotation performance.** Bars show gene-level F1 scores (left y axis), and lines with circles show the numbers of predicted genes (right y axis) for *Arabidopsis thaliana*, *Argiope bruennichi* and *Homo sapiens* across increasing inference context lengths. Increasing the inference context length altered both gene-level accuracy and the number of reconstructed gene models in a species-dependent manner.

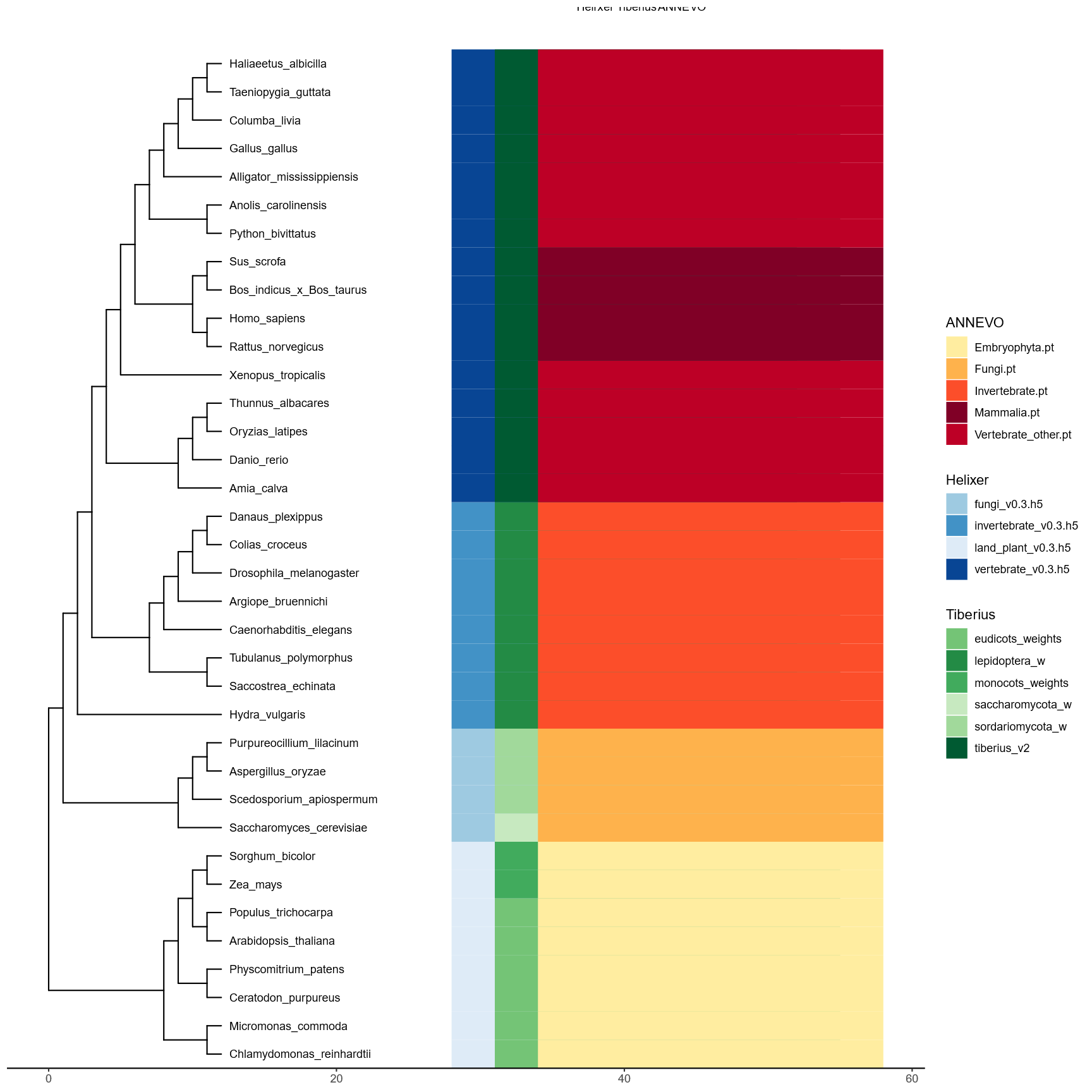

**Supplementary Fig. 11 | Method-specific pretrained model assignment for benchmark species.** The 36 benchmark species are shown according to their phylogenetic relationships (left), together with the pretrained models selected for Helixer, Tiberius and ANNEVO (right). Colored blocks indicate the model used for each species–method pair, with colors corresponding to the pretrained models listed in the legends. For each method, the model trained on the same or the nearest available taxonomic group was preferentially applied.

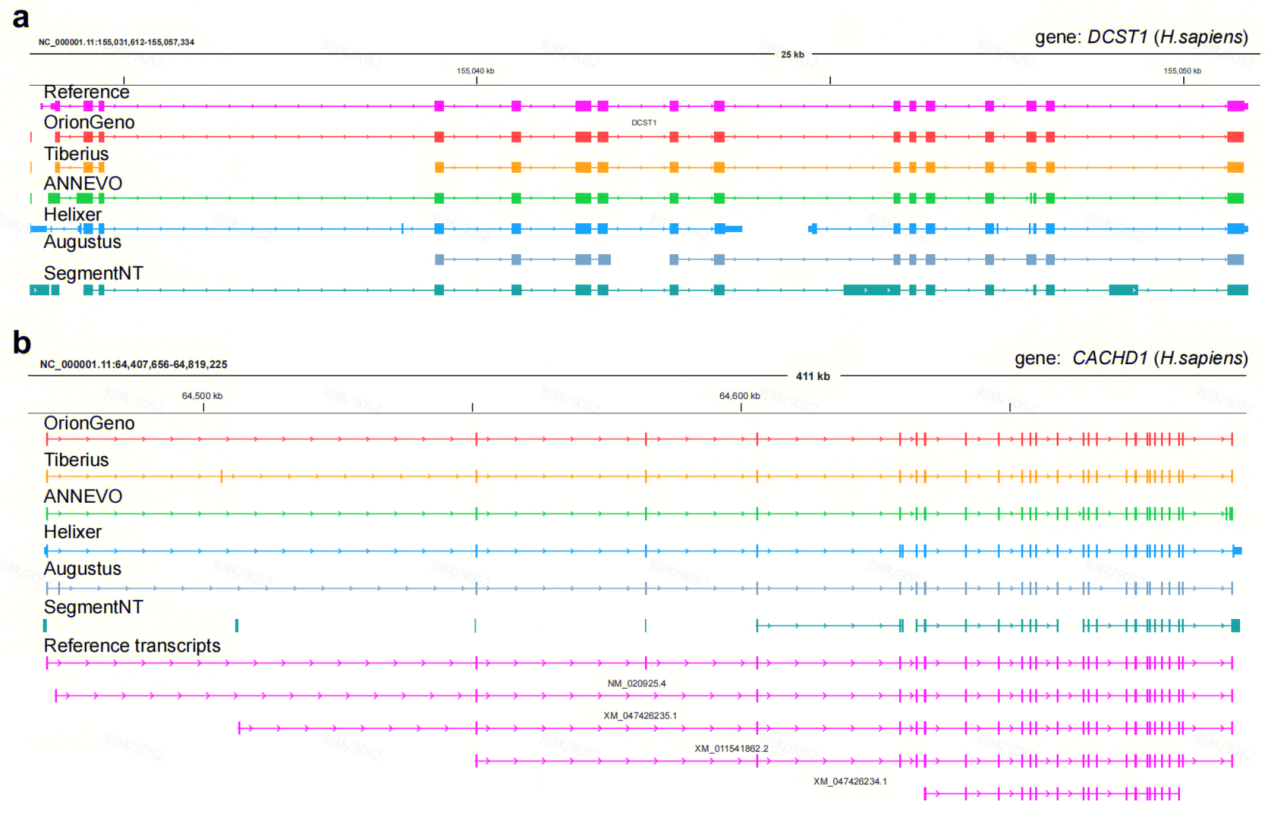

**Supplementary Fig. 12 | Comparative visualization at human *DCST1* locus (Case in Tiberius study).** SegmentNT is included by manually reconstructing its token-level probabilities into standardized GTF format. Only OrionGeno predicts the correct gene model. ANNEVO and segmentNT fail to detect the correct exons, while Tiberius, Helixer and Augustus split it into two short genes.

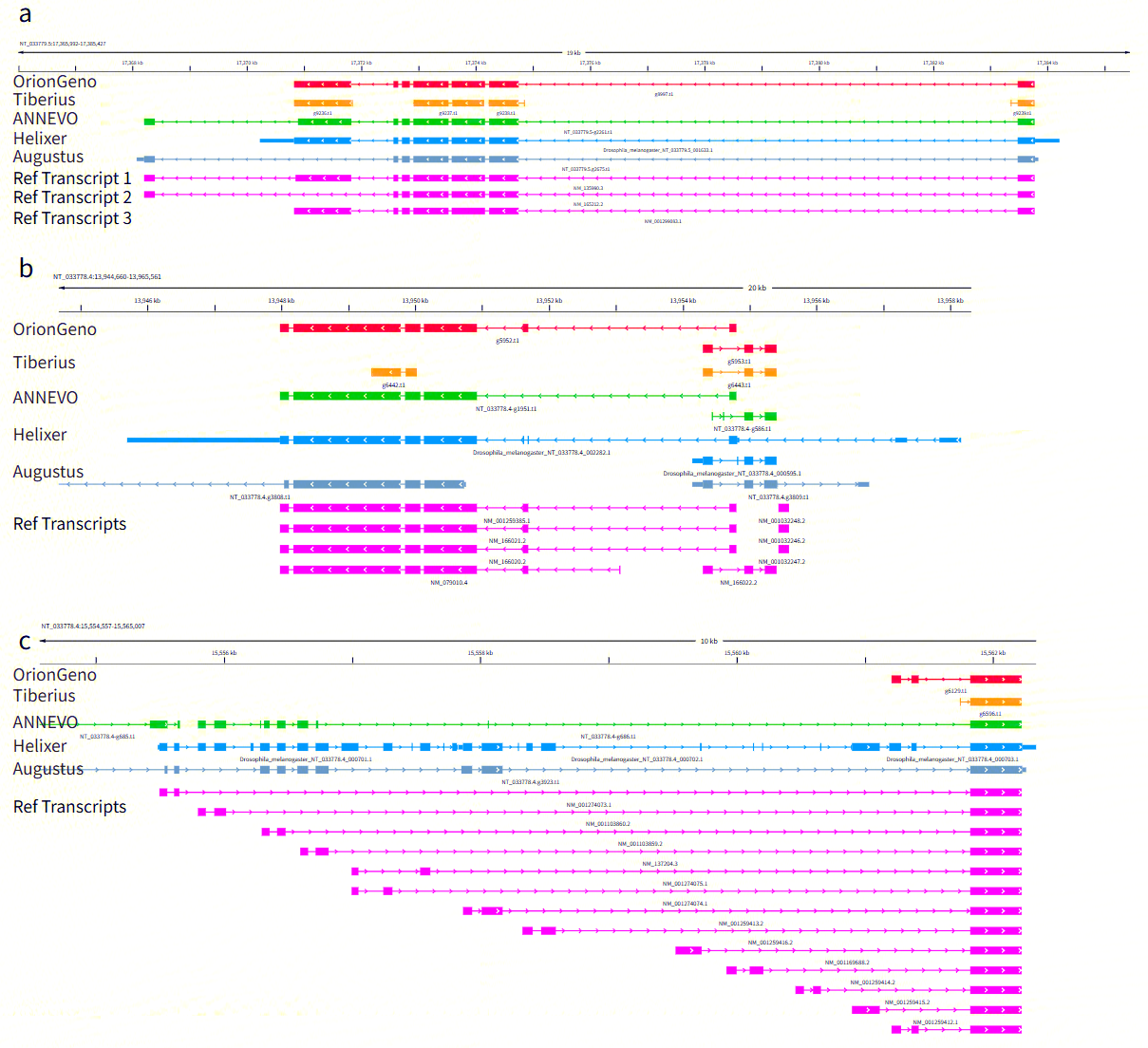

**Supplementary Fig. 13 | Comparative visualization of OrionGeno’s accurate gene prediction across example loci in *Drosophila melanogaster*.** (a–b) Representative genomic browser tracks showcasing the predicted gene structures by OrionGeno (red) and four state-of-the-art *ab initio* predictors: Tiberius, ANNEVO, Helixer, and Augustus, against ground-truth Reference Transcripts.

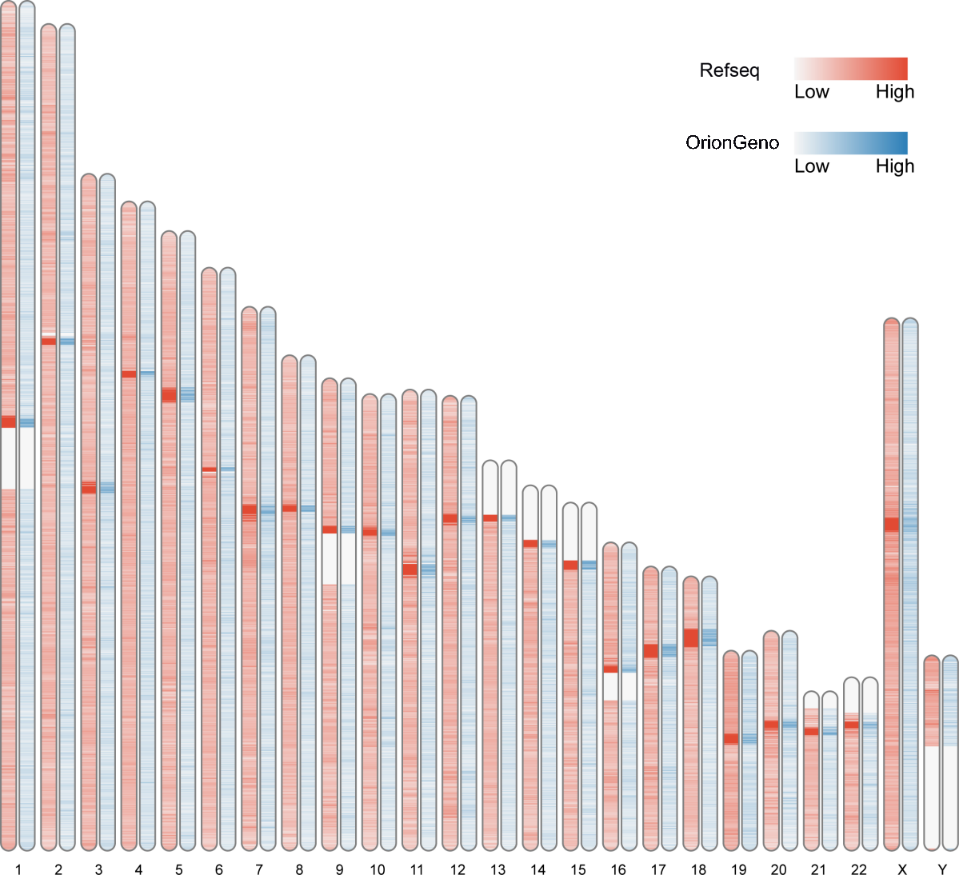

**Supplementary Fig. 14 | Global landscape of repetitive element distributions across the human genome.** Chromosome idiogram heatmap comparing repetitive element densities between the RefSeq gold-standard annotation (left tracks) and OrionGeno *ab initio* predictions (right tracks) across the human reference genome (GRCh38).
